# A feedback control for restraining autoimmune γδ T cells: reprogramming into ILC1s

**DOI:** 10.64898/2026.06.05.730476

**Authors:** S. Bajana, A. Pankow, K. Liu, N. Guzniczak, H. Bagavant, M.L. Joachims, M. Zhao, W.R. Chen, A.D Farris, U. Deshmukh, X.-H. Sun

## Abstract

γδ T cells are promising mediators of cancer immunotherapy, yet their potential to drive autoimmunity remains incompletely understood. Here, we identify a feedback mechanism in which innate-like Vγ1.1⁺Vδ6.3⁺ T cells are reprogrammed into ILC1-like cells, thereby restraining autoimmune pathology. We define a previously unrecognized ILC1 subset whose development depends on an intact *Tcrd* locus. These cells predominantly harbor productive Vγ1.1 and Vδ6 rearrangements, consistent with their origin from Vγ1.1⁺Vδ6.3⁺ T cells. Mechanistically, TCR signaling induces Id3, which suppresses E protein–dependent activation of T cell–specific genes, including that encoding Vδ6.3. Id3 ablation drives robust expansion of Vγ1.1⁺Vδ6.3⁺ T cells and severe autoimmunity, characterized by tissue infiltration, autoantibody production, enhanced T follicular helper cell differentiation, and accumulation of germinal center and age-associated B cells. Together with previously described exocrine dysfunction, these features resemble human Sjögren’s disease. Consistent with this, we observed in the salivary glands of Sjogren’s disease patients an increased frequency of CD4⁻CD8⁻ T cells enriched for γδ T cells, including subsets functionally analogous to murine Vγ1.1⁺Vδ6.3⁺ cells. Collectively, these findings uncover a TCR–Id3–dependent reprogramming pathway that limit the pathogenic potential of harmful γδ T cells.

## Introduction

The defining feature of T lineage cells is the T cell receptor (TCR) expressed on their surface. In addition to αβ T cells, γδ T cells form a critical branch of adaptive immunity by modulating immune responses in both lymphoid organs and peripheral tissues^1, 2, 3^. TCR signaling strength plays a crucial role in determining the fate between αβ and γδ T cells, as well as in the differentiation of γδ T cell subsets. Generally, γδ T cells are thought to experience stronger TCR signaling than αβ T cells^4^. Although less well understood compared to αβ T cells, the strength of TCR signaling is key to γδ T cell survival and depletion. However, whether continuous TCR signaling is required to maintain γδ T cell identity has not yet been well documented.

One of the major downstream events of γδ TCR signaling is the upregulation of the *Id3* gene, which encodes an inhibitor of E proteins, a family of basic helix-loop-helix transcription factors^4, 5^. E proteins are known to activate genes essential for T cell development, such as *Notch1* and *Cd3*^6, 7, 8^. Recently, binding sites for E proteins have also been identified upstream of a gene encoding a subunit of TCRδ, *Vδ6,* which are instrumental for the generation of Vδ6.3^+^ cells^9^. Conversely, in Id3 deficient mice, Vδ6.3^+^ T cells are dramatically expanded in ^10, 11^. These findings thus underscore the importance of the E-Id axis in regulating γδ T cell homeostasis.

γδ T cells as part of the unconventional T cells represent a non-redundant population that unlike most αβ T cells can acquire effector functions during the development in the thymus. These cells can be categorized based on their effector functions: IFNγ-producing γδ1 cells, IL-17-secreting γδ17 cells, and innate-like γδ T cells, which produce IL-4, IFNγ or IL-17^12, 13, 14^. The ontogeny of γδ T cells occurs in developmental waves, from embryogenesis through adulthood. Distinct TCRγ variable regions are expressed at different stages. For instance, *Vγ3* and *Vγ4* (Garman nomenclature; *Vγ5* and *Vγ6* in Tonegawa nomenclature) are expressed during embryogenesis, while *Vγ1.1* and *Vγ2* (*Vγ1* and *Vγ4* in Tonegawa) emerge in late embryogenesis and remain predominant in adulthood. Some γδ T cell subsets are tissue-restricted-for example, DETC cells in the skin-while others are more widely distributed but exhibit tissue preferences^15, 16^. A notable subset is *Vγ1.1Vδ6.3* γδ T cells, considered innate-like^14^. Some of these cells are also referred to as γδ NKT cells due to their expression of PLZF, a transcription factor characteristic of NKT cells^11, 17^. Because they express both IFNγ and/or IL-17, these cells are pro-inflammatory, and mechanisms must exist to tightly regulate their numbers in the body. The equivalent population in humans is thought to be Vγ9Vδ2 cells, whose utility as mediators of cancer immunotherapy is being actively pursued but their auto-immunogenic potential is not known^18, 19^.

Within the spectrum of adaptive to innate immunity, γδ T cells occupy an intermediate position. While tissue residents γδ T cells display adaptive functions during inflammation or pathogenic infections, they can also sense innate signals often presented by non-MHC molecules and play a role in homeostasis or under stress^2, 15^. Adjacent to these γδ T cells are the three groups of innate lymphoid cells (ILCs), which lack antigen receptors but perform functions analogous to T cells, such as cytokine production^20, 21^. For example, Group 1 innate lymphoid cells (ILC1s) express the signature transcription factor T-bet and secrete IFNγ in response to inflammatory cytokines, including IL-12 and IL-18. Generally, ILC1s are characterized as Lin⁻CD127⁺T-bet⁺; however, they comprise heterogeneous subsets defined by distinct markers across different tissues^22^.

We have previously discovered a population of ILC1-like cells enriched in the lamina propria of the small intestine that possesses intracellular (ic) CD3 but lack surface CD3 and any form of TCRs^23^. Because their numbers were reduced in athymic nude mice, we proposed that they originate in the thymus. In this study, we show that the abundance of icCD3⁺ ILC1-like cells depend on functional *Tcrd*, suggesting they may derive from γδ T cells. Next-generation sequencing of TCR gene rearrangement events revealed that these icCD3⁺ ILC1-like cells harbor high frequencies of productive *Vδ6-Jδ1* and *Vγ1.1-Jγ4* rearrangements, comparable to γδ T cells. We propose that these ILC1-like cells are reprogrammed from *Vγ1.1⁺Vδ6.3⁺* T cells, likely due to high *Id3* expression induced by TCR signaling in these cells. Elevated Id3 levels could inhibit E protein-mediated transcriptional activation of the *Vδ6* gene as a negative feedback control^9^. This hypothesis is further supported by the dramatic expansion of *Vγ1.1⁺Vδ6.3⁺* T cells in mice with T cell–specific deletion of *Id3* and ectopic expression of a gain-of-function E protein mutant.

Whether these ILC1-like cells possess unique functions distinct from conventional ILC1s remains to be determined. However, their derivation from *Vγ1.1⁺Vδ6.3⁺* T cells may represent a mechanism to eliminate excess pro-inflammatory γδ T cells. This is evidenced by the autoimmune phenotypes observed in our T cell–specific *Id3*-deficient mice. Like germline *Id3* knockout mice^24^, these animals exhibit elevated levels of anti-nuclear autoantibodies and immune infiltrates in tissues such as the salivary glands, lungs and liver. These phenotypes were accompanied by augmented levels of B cells, particularly germinal center B cells and age-associated B cells in the spleen and/or tissues. The increased B cell activities correlated with enhanced differentiation of follicular helper cells in a CD4 T cell extrinsic manner. Together with previously reported exocrine dysfunction, these features are similar to human Sjogren’s disease. Indeed, we observed in the salivary glands of a cohort of Sjogren’s disease patients increased frequencies of CD4^−^CD8^−^ T cells, which are enriched of γδ T cells including subsets functionally analogous to murine Vγ1.1⁺Vδ6.3⁺ cells. Taken together, our findings reveal a previously unrecognized mechanism by which potentially autoreactive γδ T cells are restrained through reprogramming into ILC1-like cells. These results also highlight the pathogenic potential of Vγ1.1⁺Vδ6.3⁺ T cells and underscore the need for caution in the design of cancer immunotherapies.

## Results

### Effector Functions of icCD3^+^ ILC1-like Cells

We previously identified a population of icCD3⁺ ILC1-like cells in the lamina propria of the small intestine^23^. To further characterize the effector functions of these cells, we compared their surface phenotypes and cytokine production capabilities with those of conventional ILC1s and NK cells. Cells were isolated from the small intestinal lamina propria and stained with a cocktail of antibodies against lineage-specific markers (FcεR, B220, CD19, CD11b, Gr-1, CD11c, Ter-119, CD3ɛ, CD4, CD8α and CD49b) (Fig. 1A). Additional staining was performed using antibodies conjugated to various fluorophores against CD45, TCRβ, TCRδ, NK1.1, CD127, Thy1.2, and KLRG1, followed by intracellular staining for CD3ε and T-bet. Subsets of αβ and γδ T cells, as well as NK cells, were defined based on TCRβ, TCRδ, and NK1.1 expression (Figure 1A).

**Figure 1.**
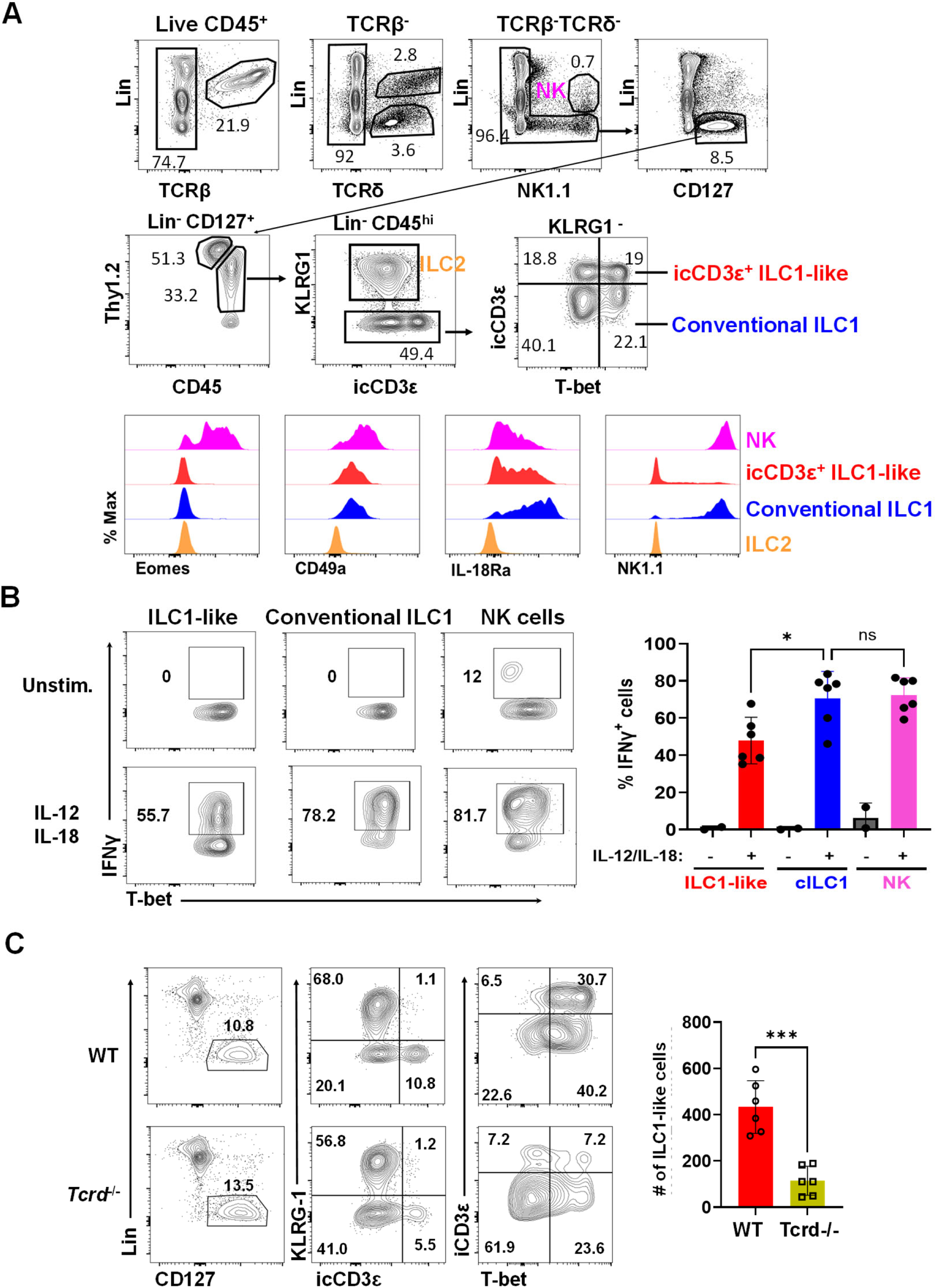
Intracellular CD3^+^ ILC1-like cells exhibit type 1 immunity and their generation is *Tcrd*-dependence. **a,** FACS analyses for ILC1 markers in cells isolated from the lamina propria of small intestine. Cell subsets of interest are labeled next to their gates. **b,** IFNγ production by indicated cell types in response to IL-12 and IL-18. One-way ANOVA was used for statistical analysis. * *p*< 0.05. **c,** Numbers of icCD3^+^ ILC1-like cells in the lamina propria of the small intestine of WT and *Tcrd*^−/−^ mice. Mann-Whitney test was used to determine the statistical significance ** *p*< 0.01.

To analyze ILCs, the population lacking αβ and γδ T cells as well as NK cells was gated as Lin⁻CD127⁺, which encompasses all ILC subsets (Fig. 1A). CD45 expression intensity was used to distinguish ILC3s (CD45^lo^) from CD45^hi^ subsets, the latter including ILC2s (KLRG1⁺) and ILC1s. Intracellular staining for T-bet and CD3ɛ allowed further discrimination between conventional ILC1s and icCD3⁺ ILC1-like cells. Additional surface markers associated with type 1 immune responses were assessed across NK cells, conventional ILC1s, and icCD3⁺ ILC1-like cells, with ILC2s serving as a negative control (Figure 1A). As expected, NK cells expressed the transcription factor EOMES, while both conventional and icCD3⁺ ILC1s did not. NK cells also expressed higher levels of CD49a than either ILC1 subset. However, icCD3⁺ ILC1-like cells exhibited lower IL-18Rα expression and lacked NK1.1, distinguishing them from conventional ILC1s.

To assess the functional capacity of icCD3⁺ ILC1-like cells, 2-month-old C57BL/6 mice were injected with IL-12 and IL-18 in the presence of brefeldin A. Cells were harvested from the lamina propria of small intestine three hours post-injection and subjected to intracellular staining for IFNγ (Figure 1B). Treatment with these pro-inflammatory cytokines robustly induced IFNγ production in NK cells, conventional ILC1s, and icCD3⁺ ILC1-like cells. Although IFNγ expression was somewhat lower in icCD3⁺ ILC1-like cells compared to the other two populations, these results clearly demonstrate their capacity to mount a type 1 immune response.

### The Generation of icCD3⁺ ILC1-like Cells Depends on Functional TCRδ

To investigate the role of TCR signaling in the differentiation of ILC1-like cells, we analyzed their presence in TCR-deficient mice. Unexpectedly, the number of ILC1-like cells in the small intestine was significantly reduced in *Tcrd*⁻^/^⁻ mice, whereas their levels were not diminished in *Tcrb*⁻^/^⁻ mice (Figure 1C, data not shown). These results suggest that the development of ILC1-like cells specifically depends on functional TCRδ, even though these cells do not express TCRδ on their surface. To further examine this, we performed intracellular staining for TCRδ in ILC1-like cells and compared them with surface TCRδ⁺ T cells and ILC2s. The staining intensity in ILC1-like cells was comparable to that in ILC2s (used as a negative control), and markedly lower than in T cells (Figure S1A).

To determine whether ILC1-like cells might be γδ T cells that had transiently downregulated surface TCRs, we cultured Lin⁻CD45^hi^KLRG1⁻NK1.1⁻ cells (a population enriched for ILC1-like cells) in the presence of IL-2 and IL-7 for three days. Very few cells re-expressed TCRδ (Figure S1B). In contrast, γδ T cells cultured under the same conditions retained TCRδ expression on their surface, arguing against transient TCR loss in ILC1-like cells.

### Distinct Transcriptional Profiles Between ILC1-like and γδ T Cells

To compare transcriptional signatures between ILC1-like and γδ T cells, we performed single-cell RNA sequencing on Lin⁻TCRδ⁺ T cells and Lin⁻CD127⁺CD45^hi^KLRG1⁻ cells (encompassing both conventional ILC1 and icCD3⁺ ILC1-like populations). We excluded Lin^+^TCRδ^+^ cells in the study because they were likely contaminating intraepithelial γδ T cells which express CD8 (Figure S2). Data analysis revealed 15 clusters found in different proportions in the two cell preparations (Figure 2A). While γδ T cells broadly expressed *Cd3g*, expression in ILC1-like cells was largely restricted to clusters 3, 5, and 6 (Fig. 2B). These three clusters were further reanalyzed and subdivided into six sub-clusters (Figure 2C). Sub-cluster 5 comprises very few cells and was then excluded in the subsequent analysis. Dot plot analysis showed that sub-clusters 1 and 2 in ILCs had over 50% of cells expressing *Cd3e* or *Cd3g*, suggesting that these sub-clusters correspond to icCD3⁺ ILC1-like cells (Figure 2D). However, their expression levels were significantly lower than those observed in T cells except *Cd3g* in sub-cluster 2 where expression in ILCs was higher but in smaller fraction of cells. Similarly, expression of *Trdc*, encoding the constant region of *Tcrd* was also lower in ILC1-like cells, potentially reflecting reduced *Tcrd* transcription.

**Figure 2.**
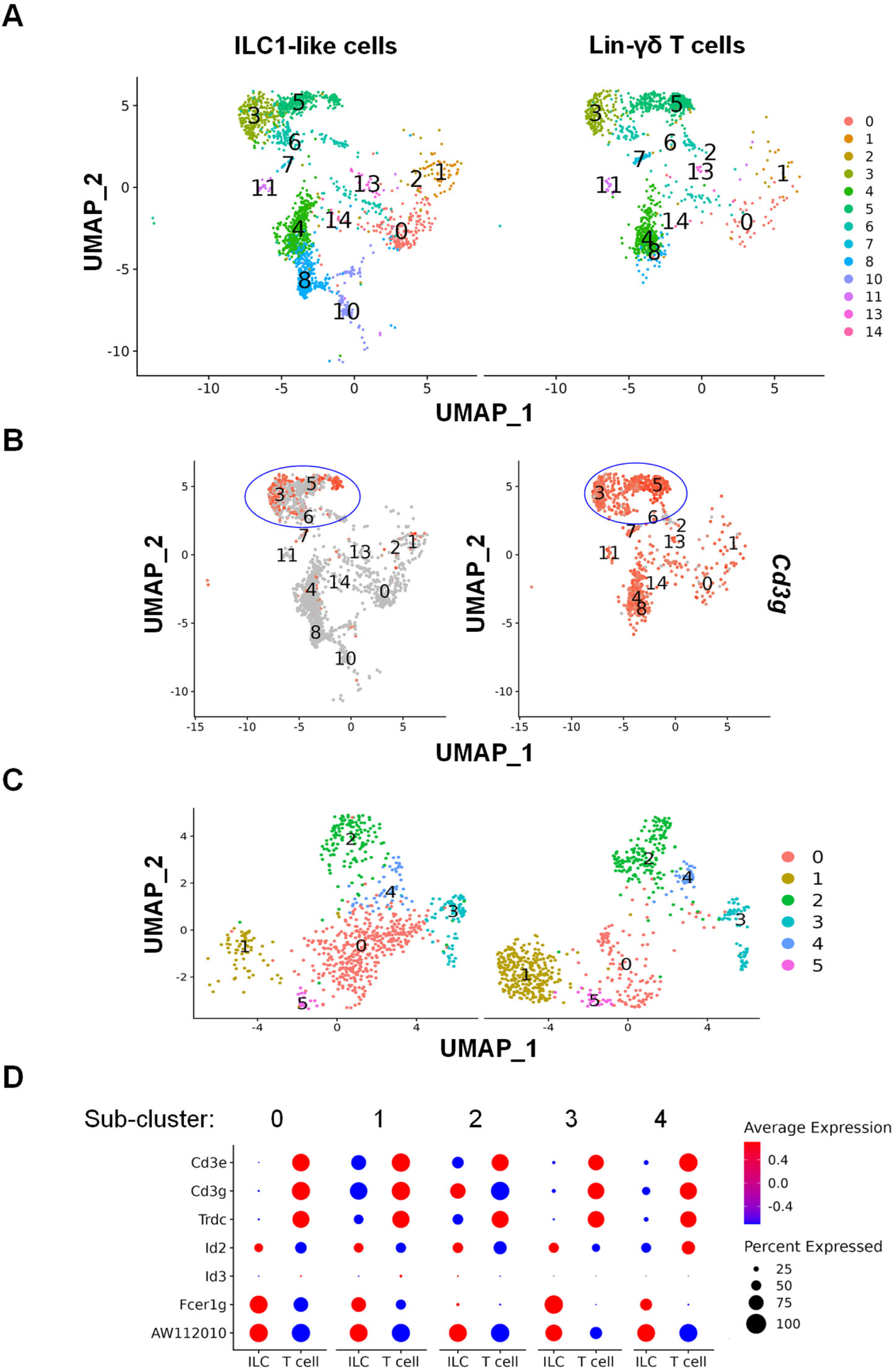
Single-cell transcriptomes of γδ T cells and ILCs. Lin^−^TCRδ^+^ T cells and Lin^−^CD127^+^CD45^hi^ KLRG1^−^ cells (ILCs) were isolated from the lamina propria of the small intestine of 2-month-old C57BL/6 mice. **a,** UMAPs of the indicated cells. Clusters are numbered and color-coded. **b,** Feature plots for the level of *Cd3g*. **c,** Sub-clustering of clusters 3, 5 and 6. **d,** Dot plots show the levels of the indicated genes and the percentages of cells in the cluster expressing the genes.

Sub-clusters 0, 3, and 4 did not show *Cd3 and Trdc* expression and are likely conventional ILC1s (Figure 2D). These sub-clusters, along with sub-clusters 1 and 2 of the ILC population, showed high expression of *Fcer1g* and *AW112010*, which were largely absent or expressed at lower levels in T cells. This pattern suggests transcriptional similarities between conventional and CD3⁺ ILC1-like cells.

Given the critical roles of Id2 and Id3 in ILC differentiation and T cell development, we assessed their expression across sub-clusters. Id2 levels were higher in ILC clusters compared to T cells, while Id3 was not detected in any sub-cluster (Figure 2D). It is therefore important to consider the redundant function of Id proteins in peripheral tissues.

### ILC1-like cells are enriched of cells with productive *Vδ6-Jδ1* and *Vγ1.1-Jγ4* rearrangements

The requirement of functional TCRδ for the formation of ILC1-like cells suggests that these cells may originate from γδ T cells, and the loss of TCR allow former γδ T cells to differentiate along the ILC1 lineage. If this hypothesis is correct, we would expect to detect productive *Tcrd* and *Tcrg* rearrangements in the DNA of ILC1-like cells, even though these genes are not actively transcribed.

The complex genomic structures of *Tcrg* and *Tcrd* are schematically illustrated in Figure 4A. *Tcrg* undergoes V to J rearrangement, whereas *Tcrd* recombines through D to J followed by V to DJ rearrangement. The junctional sequence, known as CDR3 (complementarity-determining region 3), varies with each recombination event, providing the diversity necessary for TCR specificity. Using primer pairs specific for distinct rearrangement events, we performed PCR on DNA of Lin^−^ γδ T cells, icCD3^+^ ILC1s, and icCD3^−^ ILC1s isolated from the lamina propria of the small intestine of 2-month-old C57BL/6 mice. PCR products were analyzed using gel electrophoresis (Figure 3B). As expected, recombination events typically occurring during fetal development, such as Vγ3-Jγ1 and Vγ4-Jγ1, were detected at very low levels. Other rearrangements were detected not only in T cells but also in icCD3^+^ ILC1s. However, these rearrangement events were undetectable in icCD3^−^ ILC1s sorted from the same mice (Figure 3B).

**Figure 3.**
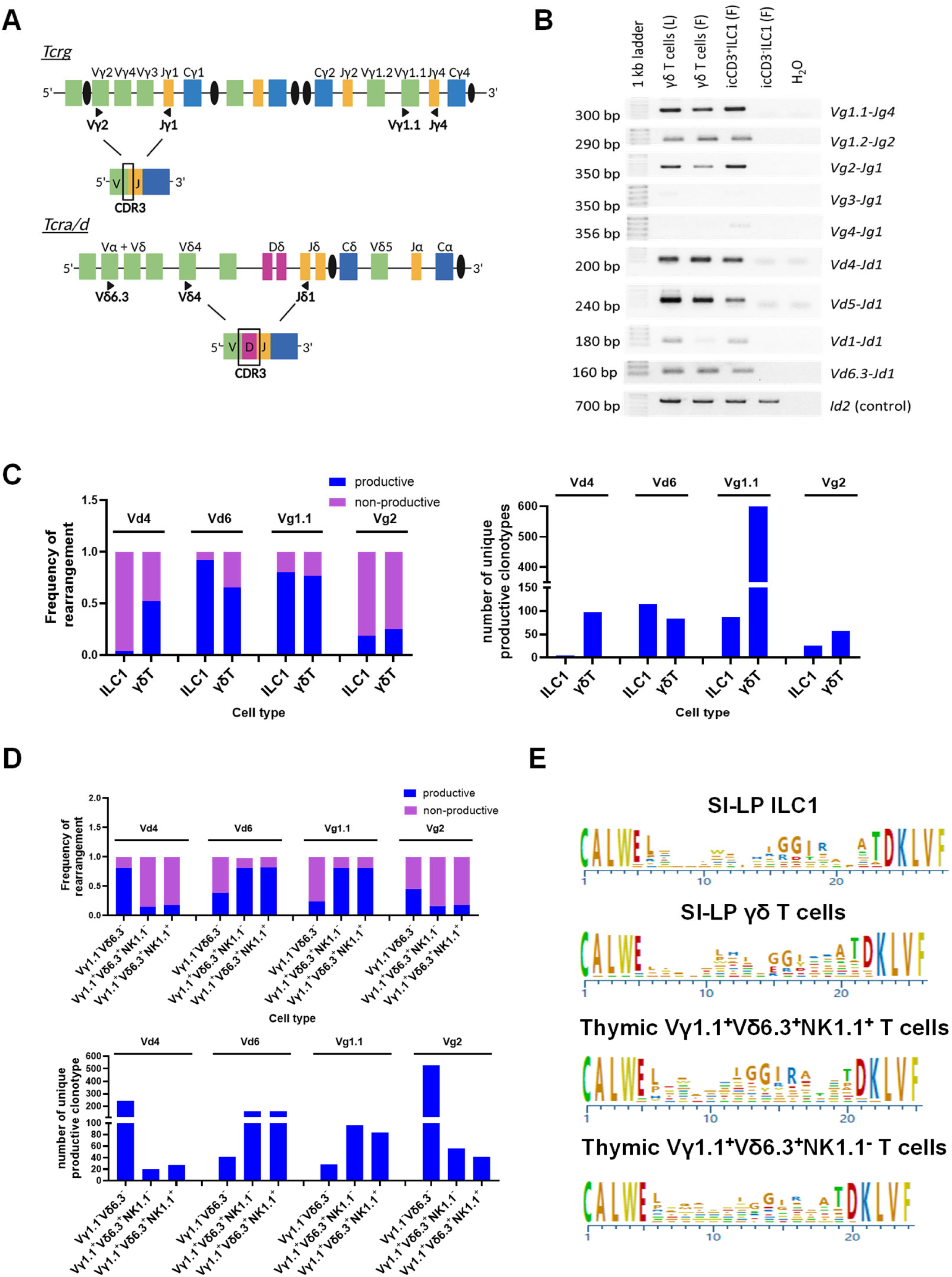
Analyses of the rearrangement of *Tcrd* and *Tcrg* show that ILC1-like cells are likely derived from Vγ1.1⁺Vδ6.3⁺ cells in C57BL/6 mice. **a,** Schematic diagrams of *Tcrd* and *Tcrg* loci. **b,** Gel electrophoretic analyses of PCR products of indicated rearrangement events with indicated DNA templates shown on top of the lanes. Live (L) or fixed (F) cells were used for isolating DNA. **c,** Frequencies of productive and non-productive clonotypes determined using deep sequencing for indicted recombination events in Lin^−^ γδ T cells and icCD3^+^ ILC1-like cells (ILC1) in the small intestine (left). The numbers of unique productive clonotypes are shown on the right. **d,** Frequencies of productive and non-productive clonotypes and the numbers of productive clonotypes for indicated rearrangement events in indicated populations of thymocytes. **e,** The consensus CDR3 sequences of the top 50 productive clonotypes in indicated cell populations.

**Figure 4.**
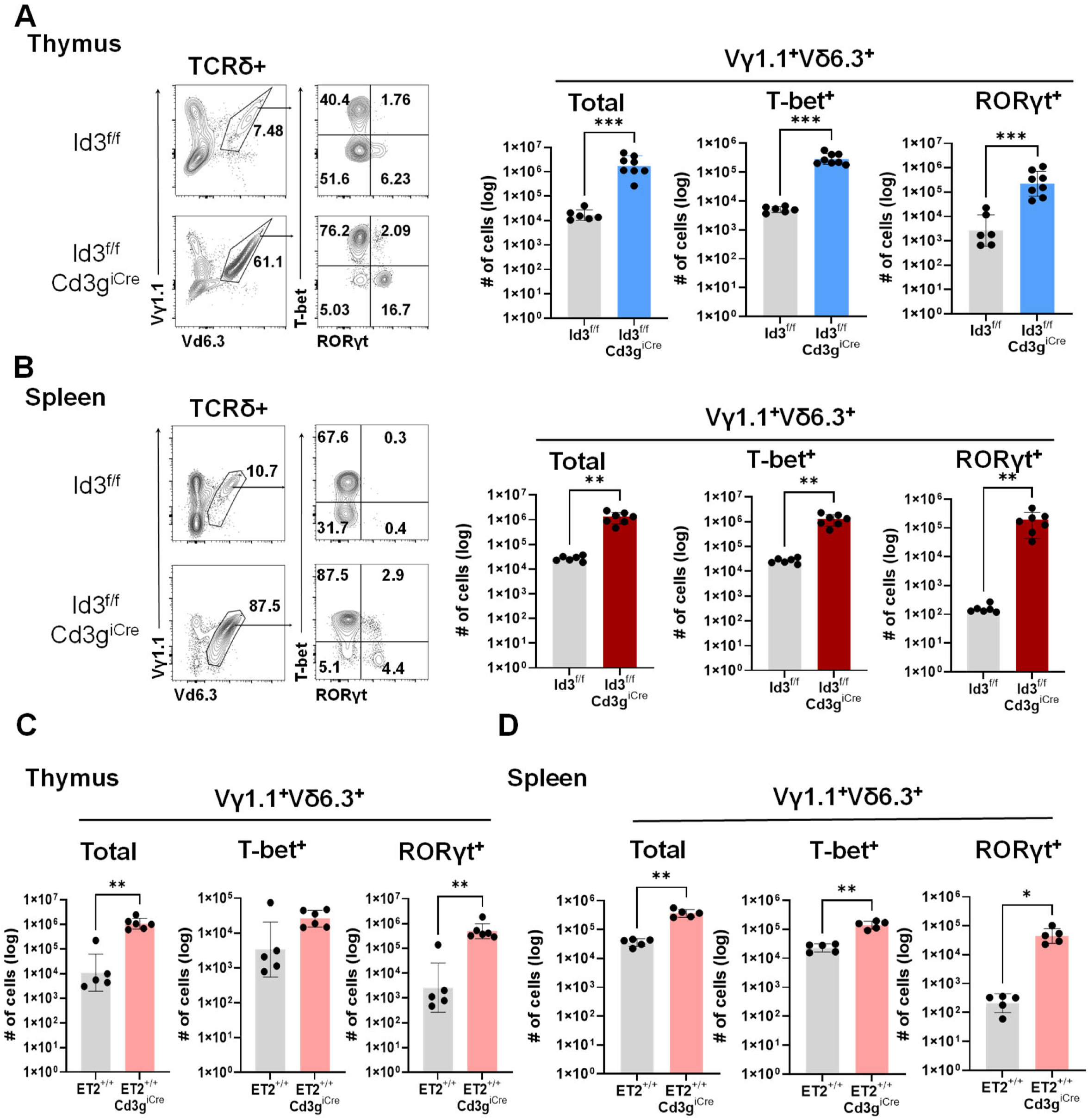
E-Id protein axis controls the abundance of Vγ1.1⁺Vδ6.3⁺ cells. Analysis of Vγ1.1⁺Vδ6.3⁺ cells in the thymus (**a**) and spleen (**b**), of 3 to 4-month-old Id3-deficient and control mice. The numbers of total and subsets of Vγ1.1⁺Vδ6.3⁺ cells are shown. The numbers of total and subsets of Vγ1.1⁺Vδ6.3⁺ cells in the thymus (**c**) and spleen (**d**) of 3 to 4-month-old *Rosa26^stop-ET^*^2^*;Cd3g^iCre^*and control mice. Mann-Whitney test was used for statistical analyses. * *p*< .05, ** *p*<0.01 and *** *p*< 0.001.

To assess the frequencies of in-frame rearrangements and the numbers of productive clonotypes, we selected representative rearrangement events in icCD3^+^ ILC1s and Lin^−^γδ T cells: Vδ4-Jδ1 and Vδ6-Jδ1 for *Tcrd,* and Vγ1.1-Jγ4 and Vγ2-Jγ1 for *Tcrg*. These samples underwent deep sequencing using an Illumina platform, generating approximately 10-40 million reads per PCR product. Clonotypes representing at least 1 in 10,000 reads were analyzed further. In the small intestine, γδ T cells exhibited over 70% productive rearrangement for Vδ4-Jδ1, Vδ6-Jδ1, and Vγ1.1-Jγ4, but only around 20% productive Vγ2-Jγ1 rearrangement, consistent with the relatively low proportion of Vγ2^+^ T cells detected using flow cytometry in this tissue (Figure 3C). In contrast, icCD3^+^ ILC1s showed very few productive Vδ4-Jδ1 rearrangements but were highly enriched for productive Vδ6-Jδ1 rearrangements. The frequency of productive Vγ1.1-Jγ4 rearrangements was comparable to that of γδ T cells, whereas Vγ2-Jγ1 rearrangements remained low, similar to γδ T cells (Figure 3C). These data suggest that icCD3^+^ ILC1s likely derive from Vγ1.1^+^Vδ6.3^+^ T cells.

We further analyzed Vγ1.1^+^Vδ6.3^+^ T cells from the thymus, comparing them to Vγ1.1^−^Vδ6.3^−^ cells as controls (Figure S3A). The Vγ1.1^+^Vδ6.3^+^ T population could be subdivided into NK1.1^+^ and NK1.1^−^ subsets. The NK1.1^+^ subset corresponds to an innate-like population, whereas the NK1.1^−^ subset includes γδ NKT cells, which express the characteristic transcription factor PLZF (Figure S3B). Similar to ILC1-like cells, both NK1.1^+^ and NK1.1^−^ Vγ1.1^+^Vδ6.3^+^ cells exhibited high frequencies of productive Vδ6-Jδ1 rearrangements and low frequencies of Vδ4-Jδ1 rearrangements (Figure 3D). The patterns of productive Vγ1.1-Jγ4 and Vγ2-Jγ1 rearrangements also resemble those of ILC1-like and γδ T cells in the small intestine, further supporting their shared gene rearrangement history. Conversely, Vγ1.1^−^Vδ6.3^−^ cells, predominantly Vγ2^+^, displayed higher frequencies of productive Vδ4-Jδ1 than Vδ6-Jδ1 rearrangements, as well as greater Vγ2-Jγ1 than Vγ1.1-Jγ4 recombination (Figure 3D).

Using the top 50 clonotypes from each cell subset, we generated consensus CDR3 sequences resulting from Vδ6-Jδ1 rearrangements (Figure 3E). Notably, the CDR3 sequences of ILC1-like cells in the small intestine closely resembled those of thymic NK1.1^+^Vγ1.1^+^Vδ6.3^+^ cells, whereas small intestine γδ T cells showed CDR3s more similar to thymic NK1.1^−^Vγ1.1^+^Vδ6.3^+^ cells. These findings suggest that ILC1-like cells originate from conventional Vγ1.1^+^Vδ6.3^+^ cells, likely through downregulation of *Tcrd* expression in the thymus. Additionally, the CDR3 sequences from these adult subsets exhibited extensive nucleotide additions and deletions, in contrast to fetal Vδ6.3^+^ cells, which have limited diversity due to the absence of terminal deoxynucleotidyl transferase^25, 26, 27^.

### The E-Id Protein Axis Regulates the Abundance of Vδ6.3⁺Vγ1.1⁺ Cells

Stimulation through the Vγ1.1/Vδ6.3 TCR induces strong signaling events, including transcriptional upregulation of Id3, which encodes an inhibitor of E protein transcription factors such as E2A and HEB (also known as TCF3 and TCF12)^4, 28, 29^. E proteins are critical for early T cell development by promoting expression of genes such as Notch1 and CD3^7^. Notably, two E protein binding sites have been identified near the *Trdv6* gene segment (encoding Vδ6.3), and these sites are essential for the generation of Vγ1.1⁺Vδ6.3⁺ cells^9^. In line with this, genetic ablation of *Id3* leads to a marked expansion of this subset^30, 31^. To investigate this in a lineage-specific manner, we employed our newly generated *Cd3g*^iCre^ transgene to delete *Id3* specifically in T cells. This resulted in a 142-fold and 48-fold increase in Vγ1.1⁺Vδ6.3⁺ cells in the thymus and spleen, respectively, compared to littermate controls lacking the *Cd3g*^iCre^ gene (Figure 4A and B).

If ILC1-like cells arise from Vγ1.1⁺Vδ6.3⁺ cells via TCR downregulation mediated by Id3, one might predict that Id3 deficiency would reduce the generation of ILC1-like cells. However, Id2, a homolog of Id3, is also expressed in these cells and may compensate for Id3 loss (Figure 2D). Accordingly, we did not observe significant changes in ILC1-like cell numbers in the small intestine of Id3-deficient mice but detected a 41-fold expansion of Vγ1.1⁺Vδ6.3⁺ cells (Figure S4A and S4B). Given the dramatic increase in Vγ1.1⁺Vδ6.3⁺ cells, a modest rise in ILC1-like cells could even occur in Id3-deficient mice if Id2 levels are high enough to have compensatory effects.

To further test whether Id3 regulates Vγ1.1⁺Vδ6.3⁺ cell numbers by repressing E protein activity, we employed a gain-of-function approach. A ROSA26 knock-in mouse strain, *Rosa26^Stop-^*ET2, expresses a mutant form of E2A (ET2) that enhances E protein activity by competing with Id proteins to dimerize with endogenous E proteins and activate E protein target genes^32, 33^. We crossed this strain to *Cd3g*^iCre^ mice, and expression of ET2 led to a substantial increase in Vγ1.1⁺Vδ6.3⁺ cells in both thymus (185-fold) and spleen (9-fold) (Figure 4C and D). These findings support a model in which E protein activity promotes, and Id3 represses, the development of this subset.

Moreover, the expanded Vγ1.1⁺Vδ6.3⁺ cells expressed either T-bet or RORγt, but the increase of RORγt^+^Vγ1.1⁺Vδ6.3⁺ cells was more pronounced in both Id3 deficient and ET2-expressing mice. The selective enrichment of the RORγt^+^ subset would suggest an augmentation of IL-17 production. To test this idea, we purified splenic Vγ1.1⁺Vδ6.3⁺ cells were cultured for three days in anti-TCRδ antibody–coated 96-well plates in medium containing IL-2 and IL-7, and measured the levels of IL-4, IFNγ and IL-17 in the culture supernatants using multiplex cytokine assays (Figure S5A). Although wild-type cells exhibited greater induction of IL-4 and IFNγ compared with Id3-deficient cells, Id3-deficient cells produced higher levels of IL-17 whereas the wild type cells did not. Consistently, we detected significant increase in IL-17 in the serum of 7 or 8-month-old Id3-deficient mice (Figure S5B). Upon stimulation of splenocytes with PMA and ionomycin *in vitro*, Id3 deficient Vγ1.1⁺Vδ6.3⁺ cells robustly produced IL-17 and IFNγ in the RORγt^+^ and T-bet^+^ subsets, respectively (Figure S5C). Therefore, the markedly increased numbers of RORγt^+^Vγ1.1⁺Vδ6.3⁺ cells and increased IL-17 production in Id3-deficient mice may have a substantial impact on immunity.

### Enhanced Differentiation of T Follicular Helper Cells in Id3-Deficient Mice

To examine the autoimmunity in Id3-deficient mice, we measured serum levels of anti-nuclear antigen antibodies (Figure 5A). We first employed a classical immunofluorescence assay to detect nuclear staining and scored the results using established criteria^34, 35^. This analysis revealed a significant increase in anti-nuclear antibodies in Id3-deficient mice compared with controls. We next quantified antibodies against double-stranded DNA and nuclear proteins using ELISA, which confirmed elevated autoantibody levels in Id3-deficient mice (Figure 5A).

**Figure 5.**
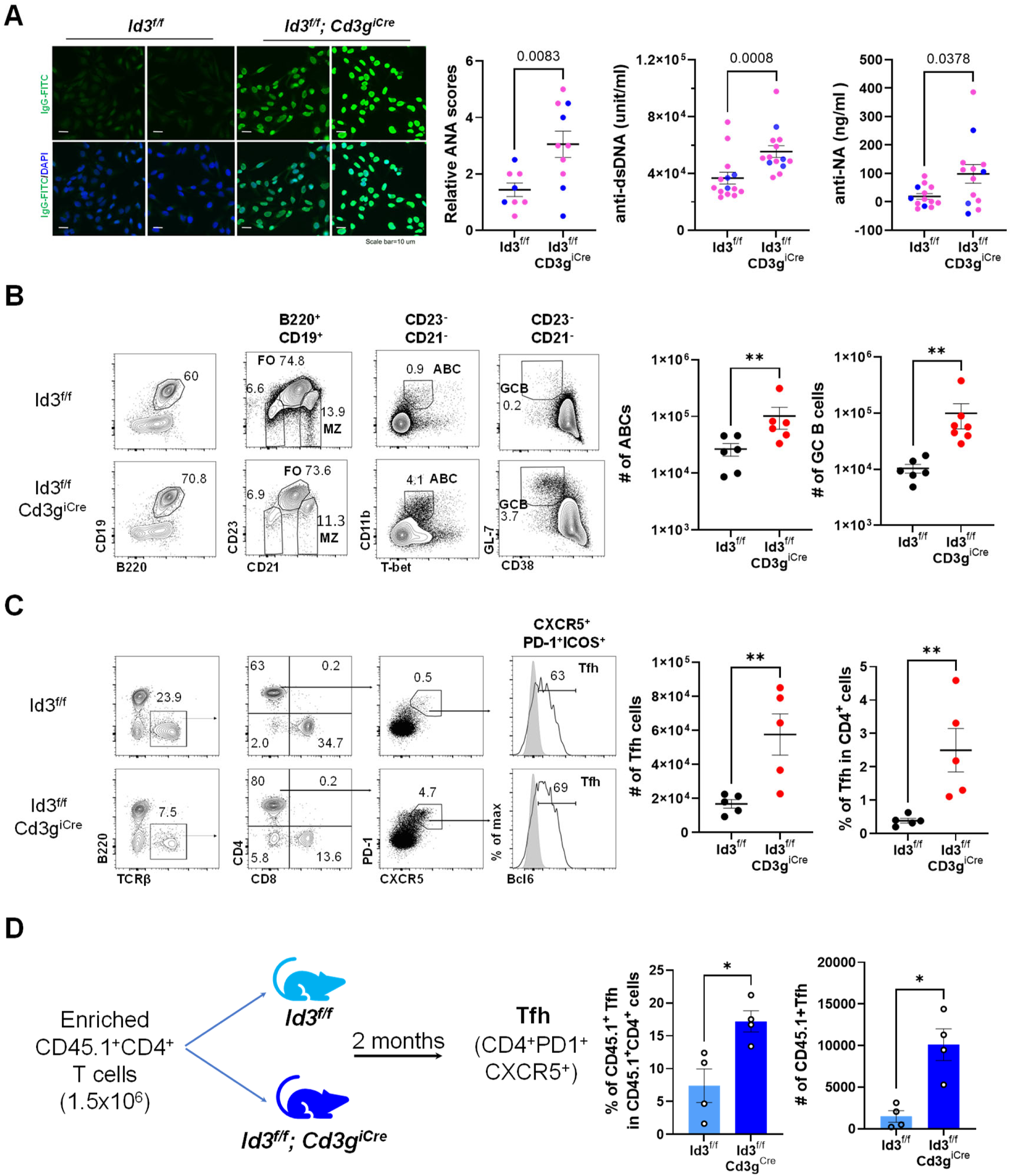
Elevation of autoantibody production and differentiation of T follicular helper cells in Id3-deficient mice. **a,** Detection of autoantibodies against nuclear antigens using immunofluorescence staining or ELISA for indicated antigens in the serum of 7-8 months old mice of the indicated genotypes. **b,** FACS of splenocytes for B cell subsets: FO, follicular; MZ, Marginal zone, ABC, aged B cells and GCB, germinal center B cells in 4-months old mice. **c,** FACS of T follicular helper cells (Tfh) in mice as described for (b). **d,** Adoptive transfer to splenocytes enriched of CD4 T cells from C57BL/6 mice into 6-weeks old recipients of the indicated genotypes. Tfh cells were analyzed 2 months later.

To assess whether the increased autoantibody production correlated with altered B cell activity in the spleen, we analyzed two B cell subsets critical for antibody secretion: age-associated B cells (ABC) and germinal center B cells (Figure 5B). Both subsets reside within the CD21⁻CD23⁻ compartment and were further defined as CD11b⁺T-bet⁺ for ABCs and GL7⁺CD38⁻ for germinal center B cells^36, 37^. In Id3-deficient mice, the frequencies of both populations were significantly increased relative to controls, providing a potential explanation for the elevated serum autoantibody levels.

T follicular helper (Tfh) cells play a central role in B-cell maturation and function. These cells, defined as CD4⁺CXCR5⁺PD-1⁺BCL6⁺, promote B cell activation, proliferation, and germinal center formation, as well as affinity maturation^38^. Consistent with the increased B cell functions observed, Id3-deficient mice exhibited a marked expansion of splenic Tfh cells in steady state, which likely contributes to enhanced B cell activity and autoantibody production (Figure 5C).

Rezende *et al*. previously reported that γδ T cells can promote Tfh differentiation in wild-type mice^39^. It is therefore plausible that the marked expansion of Vγ1.1⁺Vδ6.3⁺ γδ T cells in Id3-deficient mice exerts a similar effect. However, because Id3 regulates multiple aspects of T cell biology, we sought to exclude the possibility that increased Tfh differentiation resulted from intrinsic Id3 deficiency in CD4⁺ αβ T cells. To address this, we performed adoptive transfer experiments in which CD45.1⁺ splenocytes enriched for CD4⁺ T cells were transferred into Id3^f/f^ mice with or without *Cd3g^iCre^*. Two months after transfer, donor T cell differentiation was assessed. Both the frequency and absolute number of donor-derived Tfh cells were significantly higher in Id3-deficient recipients than in controls, indicating that Id3 deficiency creates an environment conducive for Tfh differentiation, most likely mediated by the expanded Vγ1.1⁺Vδ6.3⁺ γδ T cell population (Figure 5D).

### Autoimmune Phenotypes in the Salivary Glands and Lung of Id3-Deficient Mice

Histological examination of the salivary glands and lungs from 7 to 8-month-old Id3-deficient mice and their littermate controls revealed pronounced immune cell infiltration in these tissues. A greater proportion of Id3-deficient mice exhibited inflammatory foci, and the number of foci per tissue section was markedly increased (Figure 6A). The salivary gland pathology closely resembles that previously reported in germline Id3 knockout mice and Lck-Cre–mediated Id3 conditional knockouts¹⁰,²², although it manifested at a markedly earlier age (7 versus 12 months old). In contrast, immune infiltration in the lung has not been previously described. Additionally, immune cell infiltration was observed in the liver of Id3-deficient mice (Figure S6A). Splenomegaly was also detected in approximately 20% of Id3-deficient mice (Figure S6B), with spleens enriched in lymphoid cells expressing T cell markers and metastasis to peripheral organs such as the lung, liver and salivary glands.

**Figure 6.**
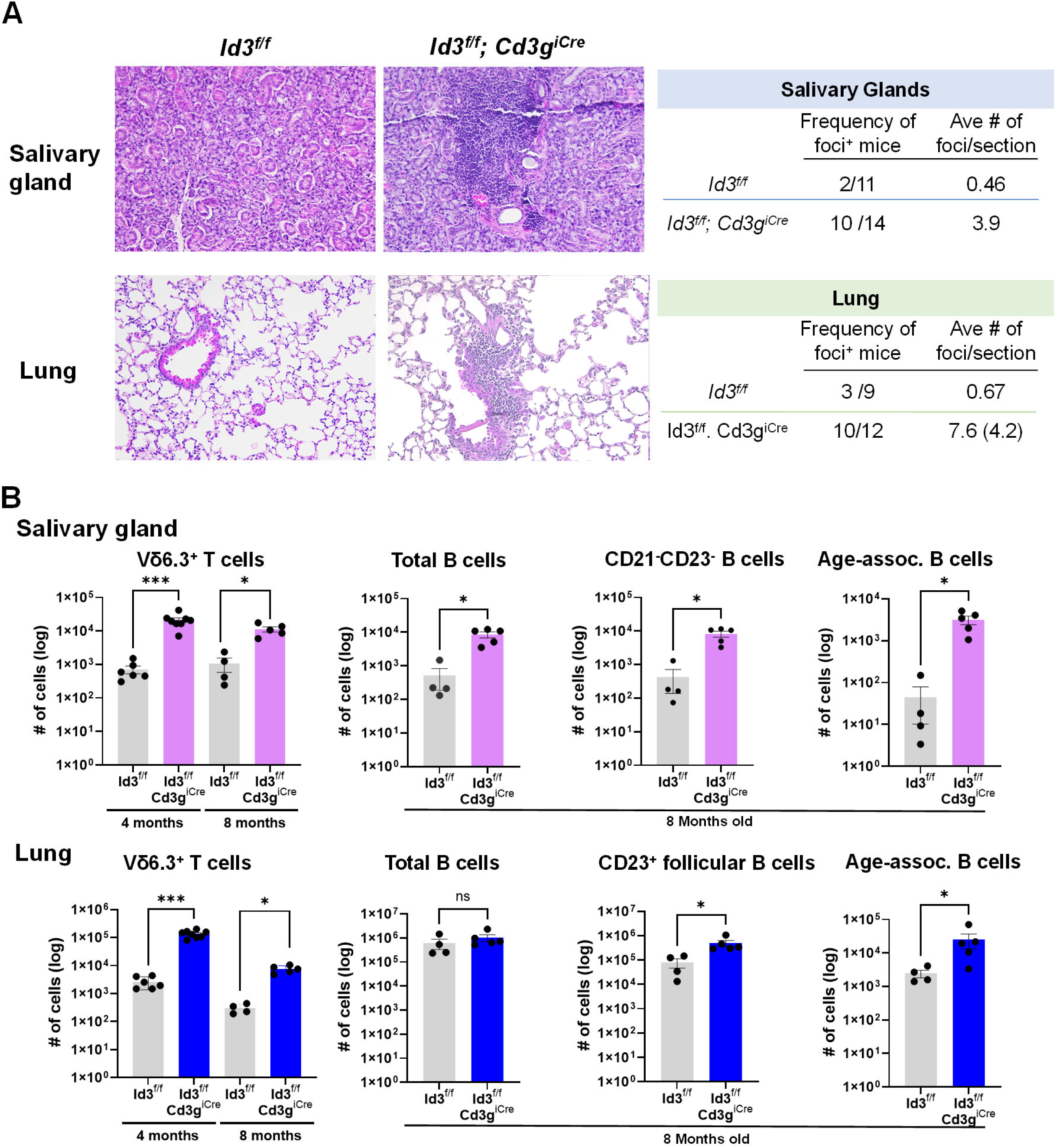
Autoimmunity developed in Id3-deficient mice. **a,** H&E staining to tissue sections from 7-8 months old mice of the indicated genotypes (left). Summaries of samples analyzed are shown on the right. **b,** Numbers of Vγ1.1⁺Vδ6.3⁺ cells and B lineage cells determined using flow cytometry in indicated tissues and ages of mice of the indicated genotypes.

Flow cytometric analysis of salivary glands and lungs from 4- and 8-month-old mice revealed a dramatic expansion of Vγ1.1⁺Vδ6.3⁺ cells at both ages (Figure 6B). In the salivary glands, the B cell compartment was profoundly altered: total B cell numbers were increased approximately 17-fold in Id3-deficient mice, primarily due to expansion of the CD23⁻CD21⁻ population, which includes age-associated B cells (ABC, CD11b⁺T-bet⁺) linked to autoimmunity^40^. Although total B cell numbers in the lung were unchanged, CD23⁺ follicular B cells and ABCs were increased by approximately 3.5-fold and 10-fold, respectively (Figure 6B). Collectively, these findings implicate B cells as major contributors to the autoimmune phenotypes observed in Id3-deficient mice, potentially driven by the expanded population of Vγ1.1⁺Vδ6.3⁺ cells throughout the body.

In contrast, T cell compartments were not expanded in Id3-deficient mice. Instead, both CD4⁺ and CD8⁺ T cell numbers were significantly reduced in the lungs (Figure S6C). The number of regulatory T cells (Tregs) was also decreased correspondingly to the numbers of CD4⁺ T cells, thus keeping the ratio of Tregs versus CD4^+^ T cells unchanged in the lung. Interestingly, very few Tregs were detected in the salivary glands of either genotype. Reduced T cell numbers in peripheral tissues correlated with impaired T cell development in the thymus and decreased T cell numbers in the spleen. Similarly, the reduced number of splenic Tregs was proportional to diminished CD4⁺ T cells (Figure S6C).

To investigate the defect in thymic T cell output, we performed BrdU labeling experiments by injecting BrdU into mice 16 hours prior to the analyses. We observed a marked reduction in BrdU incorporation at the immature CD8 single-positive (ISP) stage, a developmental stage characterized by robust proliferation in response to pre-TCR signaling (Figure S6D). This finding is consistent with the known upregulation of Id3 downstream of pre-TCR stimulation and supports a role for Id3 in T cell expansion^5^. Interestingly, despite the massive accumulation of Vγ1.1⁺Vδ6.3⁺ cells in Id3-deficient mice, these cells did not exhibit increased BrdU incorporation in the thymus, thus excluding the increased proliferation as a mechanism (Figure S6C).

### Enrichment of CD4 and CD8 double negative T cells in the salivary glands of patients with Sjogren’s disease

Focal lymphocytic infiltrates in salivary gland and lung, elevated T follicular helper cells, B cell dysregulation, and anti-nuclear antibodies observed in Id3-deficient mice are also characteristic features of Sjögren’s disease (SjD)^41^. Human Vγ9⁺Vδ2⁺ γδ T cells that functionally resemble murine Vγ1.1⁺Vδ6.3⁺ γδ T cells have been shown to primarily reside in the CD4/CD8 double negative (DN) population^42^, and examination of healthy control PBMC confirmed the predominantly DN T cell phenotype of human Vγ9⁺Vδ2⁺ γδ T cells, where they comprised about 85% of all blood Vγ9⁺Vδ2⁺ γδ T cells (Figure 7A). Therefore, we leveraged our previously collected flow cytometry dataset of human salivary gland biopsies including CD45, CD3, CD4 and CD8 markers to explore the potential relevance of our findings to SjD (Table S1). The dataset included salivary gland flow cytometry data from Ro/SS-A seropositive SjD cases (n=14), Ro/SS-A seronegative SjD cases (n=9), and non-SjD sicca syndrome controls (n=38). Salivary gland DN T cell frequencies were significantly elevated in Ro seropositive SjD cases compared to non-SjD sicca controls, consistent with the possibility that Vγ9⁺Vδ2⁺ γδ T cells may contribute to SjD pathology (Figure 7B).

**Figure 7.**
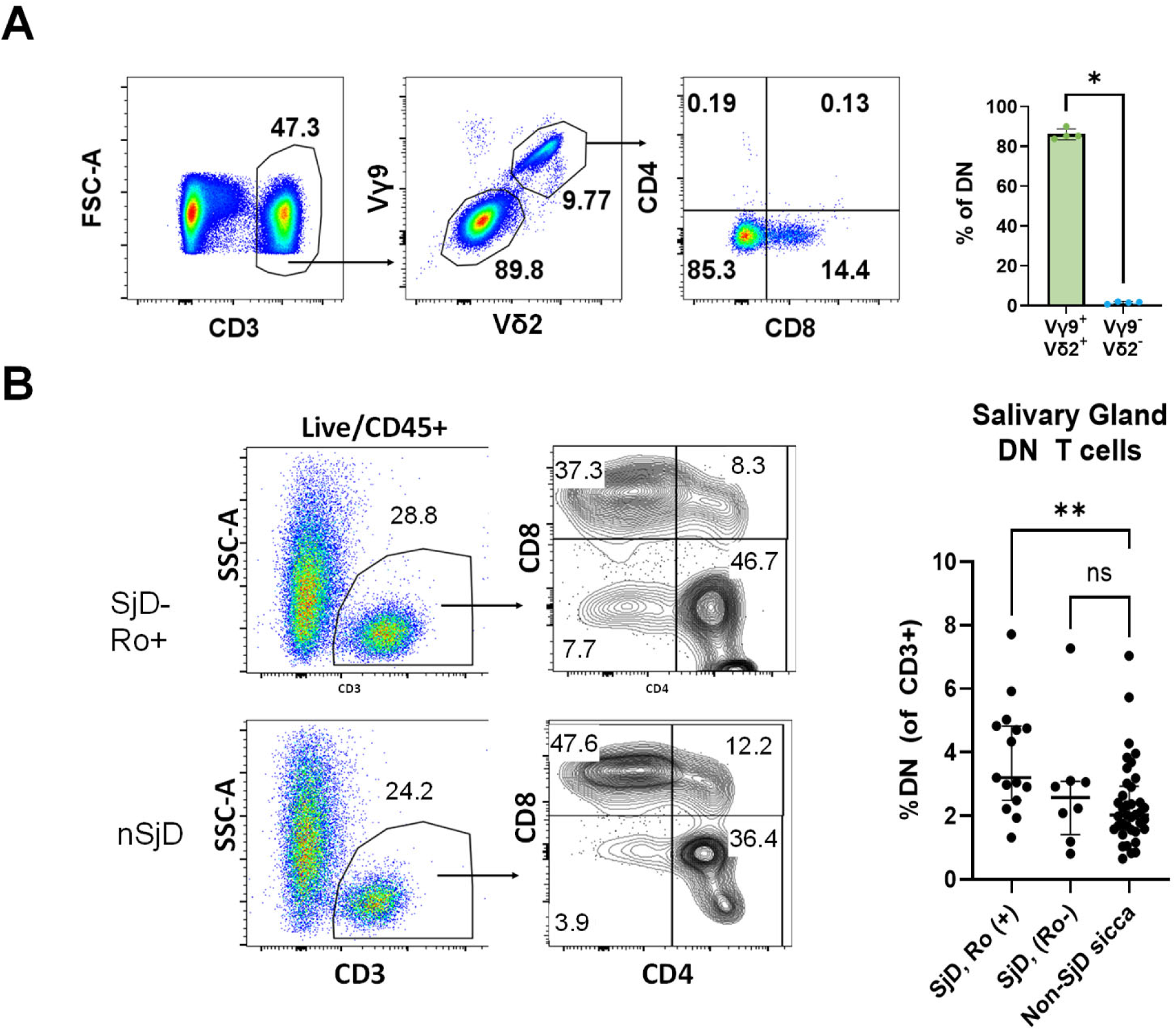
Increased frequencies of CD4 and CD8 double negative T cells in salivary glands of Sjogren’s disease patients. **a,** CD4 and CD8 profile of Vγ9^+^Vδ2^+^ γδ T cells in the PBMCs of health donors. **b,** Representative staining from SjD and nSjD-sicca patients. Comparison of CD4/CD8 DN frequency out of CD3^+^ cells between SjD-Ro+ (n=15), SjD-Ro- (n=8) and nSjD subjects (n=38). Significance determined by Kruskal-Wallis test with multiple comparisons.

## Discussion

Group 1 innate lymphoid cells comprise NK cells and ILC1s, both of which produce IFNγ in response to pro-inflammatory signals. Although these cells are thought to arise from common lymphoid progenitors in the fetal liver and bone marrow, their heterogeneity is shaped by developmental stage, activation status, tissue environment, and inflammatory cues^22^. Whether distinct subsets within this group are pre-programmed for specialized functions under specific conditions remains an open question. In this study, we add an additional layer of complexity by identifying a subset of ILC1s derived from a specific population of γδ T cells, namely Vγ1.1⁺Vδ6.3⁺ T cells. Although cellular plasticity has been described within the lineages of helper T cells or innate lymphoid cells^43, 44, 45^, our finding represents, to our knowledge, the first example of lineage conversion from an adaptive to innate cell type.

γδ T cells integrate signals from both adaptive and innate pathways, positioning them as sentinels in barrier tissues^46, 47, 48, 49^. In this context, TCR signaling is not merely required for antigen recognition but also for enforcement of T cell identity. The icCD3⁺ ILC1-like cells described here are not simply T cells that have lost surface or intracellular TCR expression. Rather, they have acquired defining features of ILC1s, including high expression of *Fcer1g* and *AW112010*, comparable to conventional ILC1s and NK cells. These findings suggest that continuous γδ TCR expression is required to maintain T-cell identity in Vγ1.1⁺Vδ6.3⁺ T cells.

The generation of CD3⁺ ILC1-like cells depends on intact *Tcrd*, indicating a requirement for a functional γδ TCR and implicating γδ T cells as their precursors. Because these cells lack both surface and intracellular TCRδ protein and display markedly reduced levels of *Trdc* transcripts, it is likely that the genes encoding TCRδ have been inactivated. To investigate their developmental history, we analyzed TCR gene rearrangements, which serve as a molecular fossil record. Deep sequencing of PCR products targeting *Tcrg* and *Tcrd* rearrangements was performed, focusing on the major γ and δ chains expressed in adult mice. Given that approximately 300 cells were used per PCR reaction and assuming at least one rearrangement per cell per gene, the large number of productive clonotypes observed indicates sufficient sequencing depth. In the small intestine, ILC1-like cells and γδ T cells exhibited similar frequencies of productive Vγ1.1–Jγ4 and Vγ2–Jγ1 rearrangements. In contrast, *Tcrd* rearrangements differed markedly between the two populations. Whereas γδ T cells showed high frequencies of both productive Vδ4–Vδ1 and Vδ6–Vδ1 recombination, ILC1-like cells predominantly harbored productive Vδ6–Vδ1 rearrangements, with very few Vδ4–Vδ1 clonotypes. These findings implicate Vγ1.1⁺Vδ6.3⁺ T cells as the origin of the ILC1-like population.

Previous studies have shown that the development of Vγ1.1^+^Vδ6.3^+^ γδ T cells in the thymus is tightly regulated by the E protein-Id3 axis^9, 30^. Engagement of the TCR by high-affinity ligands delivers strong ERK-dependent signals that induce Egr transcription factors and Id3 expression, thereby limiting transcription of *Trav15d-1-dv6d-1* gene segment encoding Vδ6.3 which is preferentially paired with Vγ1.1^9^. Id3 upregulation reduces the availability of E proteins required to bind regulatory elements within the *Trav15* locus, leading to suppression of Vδ6.3 transcription and restricting the size of this γδ T cell subset. Conversely, in the absence of Id3, elevated E-protein activity promotes increased transcription of Vδ6.3 resulting in marked expansion of the Vγ1.1^+^Vδ6.3^+^ γδ T cell compartment^30^. Our data extend this regulatory framework and suggest that Id3 not only constrains the size of this population but is also required for reprogramming of Vγ1.1^+^Vδ6.3^+^ into ILC1-like cells. Thus, we propose that strong TCR signaling engages an Egr-Id3-E protein negative feedback loop that limits the accumulation of potentially pathogenic γδ T cells and promotes their diversion into an innate-like lineage through transcriptional reprogramming toward a TCR-independent ILC1-like effector state.

Reprogramming Vγ1.1⁺Vδ6.3⁺ T cells into CD3⁺ ILC1-like cells may serve as a mechanism to restrain their inflammatory potential. Disruption of this regulatory checkpoint through Id3 deletion or enhanced E-protein activity results in massive expansion of Vγ1.1⁺Vδ6.3⁺ T cells and leads to severe autoimmunity, underscoring the importance of tightly controlling their abundance. The preferential expansion of RORγt^+^Vγ1.1⁺Vδ6.3⁺ T cells in Id3-deficient mice resulted in the elevation IL-17 production, a critical mediator of inflammatory reactions. Unlike IFN-γ which acts primarily on immune cells, IL-17 exerts its effects on non-hematopoietic cells, including epithelial and stromal compartments inducing the release of chemokines and inflammatory mediators. In this context, IL-17 promotes the recruitment of neutrophils and monocytes through induction of chemokines such as CXCL1 and CXCL2 thereby establishing a local inflammatory circuit^50^. Although autoimmune phenotypes have previously been reported in Id3-deficient mice, our use of a T-cell-specific Cre, *Cd3g^iCre^,* to ablate Id3 revealed a more widespread autoimmune pathology, affecting not only the salivary glands but also the lung and liver, along with significantly elevated autoantibody levels at younger ages than previously described^10, 24^. This phenotype is likely attributable to more efficient Id3 deletion within the γδ T-cell lineage, resulting in greater expansion of Vγ1.1⁺Vδ6.3⁺ T cells. Id3 has been shown to regulate Treg differentiation, and germline Id3 deficiency results in a reduction in Treg numbers due to repression of FoxP3 expression^51^. However, whether this reduction alone is sufficient to drive spontaneous autoimmunity remains unresolved. Miyazaki et al. demonstrated that *FoxP3^Cre^*-mediated deletion of Id3 alone did not result in autoimmune phenotypes, likely due to functional redundancy with Id2. In our T cell–specific Id3 knockout mice, we observed an approximately twofold reduction in Treg numbers accompanied by a similar decrease in total CD4⁺ T cells, resulting in comparable Treg-to-CD4⁺ T-cell ratios between Id3-deficient and control mice. The residual Treg population across multiple tissues may therefore be sufficient to restrain spontaneous autoimmune disease. However, reduced Treg numbers could potentiate autoimmunity when combined with additional abnormalities.

We further show that enhanced Tfh differentiation occurs through a CD4 T-cell-extrinsic mechanism, likely driven by the expanded Vγ1.1⁺Vδ6.3⁺ T-cell population. Wild type CD4 T cells exhibited enhanced Tfh differentiation when they were placed in Id3-deficient mice. Together with elevated local and systemic levels of IL-4, IFNγ, and IL-17, Tfh cells facilitate a variety of B cell function which leads to increased autoantibody production and B-cell infiltration.

The autoimmune phenotypes observed in *Id3*-deficient mice have been proposed to resemble Sjögren’s disease^10, 52^. Although direct equivalence between mouse models and human disease is not feasible, key features—including reduced saliva and tear production and immune infiltration of the salivary glands—mirror clinical manifestations in patients. We additionally observed immune infiltration in the lung and liver, organs frequently involved in Sjögren’s disease. Splenomegaly, which developed in a subset of Id3-deficient mice, may parallel the increased risk of lymphoma observed in some patients. Regardless of whether Id3 deficiency constitutes a faithful model of Sjögren’s disease, these findings underscore a critical role for γδ T cells in autoimmunity.

In human salivary glands, we detected a significant increase in the frequency of CD4⁻CD8⁻ (DN) T cells in Ro/SS-A seropositive Sjögren’s disease patients compared to non-Sjögren sicca controls. Like murine Vγ1.1⁺Vδ6.3⁺ T cells, human Vγ9Vδ2 T cells exhibit innate-like properties, including expression of inflammatory cytokines, PLZF and NK receptors^53, 54^. Notably, these innate-like T cells are predominantly DN, raising the possibility that they contribute to the increased DN T cell population observed in our dataset. Consistent with this notion, elevated γδ T cell frequencies have been reported in a study using human Sjögren’s disease samples^55^. Together, these observations suggest that dysregulated γδ T cells may drive systemic autoimmunity with organ-specific manifestations.

γδ T cells have also attracted considerable interest in immunotherapy due to their non-MHC-restricted activation and potent pro-inflammatory functions. These cells can be expanded *in vitro* and engineered with cancer-targeting T cell receptors. Our findings reveal previously underappreciated autoimmune-promoting properties of γδ T cells, which should be carefully considered in the design of γδ T cell–based therapeutic strategies.

## Author Contribution

S.B., A.P., M.J., N.G. and X.-H. S. performed the experiments; K.L. carried out bioinformatics analyses; S.B., A.P. H.B., M.Z. and X.-H. Sun designed the study; S.B., A.P., K.L., W.R.C. U.D., A.D.F. and X.-H. S. analyzed the data and wrote the manuscript.

## Acknowledgements

We thank Dr. Yuan Zhuang for providing Id3^f/f^ mice. We are grateful to the Clinical Genomics Center and the Flow cytometry facility at the Oklahoma Medical Research Foundation for outstanding technical assistance. This work was supported by grants from the NIH (R01 AI178947) and the RESTORE grant for American Association of Immunologists to X.-H.S., as well as the Oklahoma Center for the Advancement of Science and Technology (HR16-085) to W.R.C. The clinical research was supported by NIH grants P50AR060804 and U01DE017593. The content is solely the responsibility of the authors and does not necessarily reflect the official views of the National Institutes of Health. The Aurora flow cytometer was purchased with support from NIH grant 1S10OD028479-01. Tissue histology service provided by the tissue Pathology core was supported partly by the National Institute of General Medical Sciences Grant P30GM154635 and National Cancer Institute Grant P30CA225520 of the National Institutes of Health.

## STAR Methods

### RESOURCE AVAILABILITY

#### Lead contact

Xiao-Hong Sun, Oklahoma Medical Research Foundation, 825 NE 13^th^ Street, Oklahoma City, OK, Tel: 405-271-7103, Fax: 405-271-7128.

#### Materials availability

This study did not generate new unique reagents.

#### Data and code availability

Single cell RNAseq data has been deposited in NCBI. The accession number for the scRNA-seq data reported in this paper is GEO: **GSE317233** (https://www.ncbi.nlm.nih.gov/geo/query/acc.cgi?&acc=GSE317233).

This paper does not report original code. Any additional information required to reanalyze the data reported in this paper is available from the lead contact upon request.

Any additional information required to reanalyze the data reported in this paper is available from the lead contact upon request.

### EXPERIMENTAL MODEL AND SUBJECT DETAILS

#### Experimental model and study participant details

##### Mice

C57BL/6J mice were purchased from The Jackson Laboratory. Rosa26^stop-ET2^ (ET2) mice were generated as previously described^32^. Id3^F/F^ mice were gifts from Yuan Zhuang, Duke University^10^. *Cd3g^iCre^* mice were generated by Applied StemCell Inc. and have been described previously (Pankow and Sun). Both male and female mice were used in all experiments, and no significant sex-based differences were observed. Unless otherwise indicated, mice aged 2–4 months were used. All animal experiments were conducted in accordance with protocols approved by the Institutional Animal Care and Use Committee of the Oklahoma Medical Research Foundation.

##### Human Cohort

Participants with xerostomia and/or xerophthalmia attending the Oklahoma Sjögren’s Research Clinic between 04/01/2015 and 02/10/2016 were evaluated for fulfillment of the 2016 research classification criteria for Sjögren’s disease (SjD) as previously described ^56, 57^. Those fulfilling criteria for both SjD and other rheumatic disease(s) were excluded. Those not fulfilling SjD criteria were designated as having non-SjD sicca syndrome. Ro/SS-A autoantibody status was determined by the BioPlex™ 2200 ANA Screen test. Those having sufficient minor salivary gland biopsy tissue available for flow cytometry were included in the study.

### METHOD DETAILS

#### Cell isolation and flow cytometry analysis

Single-cell suspensions from the small-intestinal lamina propria were prepared essentially as described previously ^58^. Lungs were chopped and enzymatically digested using DNAse I (20 U/ml; Roche #10104159001) and Liberase TH (25 μg/ml; Roche #05401151001) in lung digestion buffer (10 mM HEPES, 150 mM NaCl, 5M mM KCl, 1 mM MgCl_2_, 1.8 mM CaCl2, pH 7.4) while being placed on a shaker at a speed of 210 rpm for 30-40 min at 37°C. Submandibular and sublingual salivary glands were processed by digesting the tissue in the RPMI 1640 medium (cytiva #SH30011.03) containing 40 units/ml DNase I (Roche) and 1.3 mg/ml Collagenase D (Roche #11088866001) for 1 hour at 37°C. Spleen was digested with DNAse I (27 U/ml; Roche) and Liberase TL (30 mg/ml; Roche #05401020001) for 20 min on a shaker at speed of 210 rpm in RPMI 1640 medium at 37°C. Lymph nodes were digested with DNAse I (40 U/ml; Roche) and Collagenase D (1 mg/ml; Roche) for 30 min in RPMI 1640 medium at 37°C. After the digestion, tissues were mashed with a syringe plunger on a cell strainer and washed with 1X HBSS (Gibco #14025092) with 5% newborn calf serum (Sigma #N4637). Single cell suspension from the thymus was prepared using a cell strainer and a syringe plunger.

All samples were incubated with viability dye Zombie Aqua Fixable Viability Kit (BioLegend #423101). Before adding the antibody cocktails, samples were incubated with Fc-block (CD16/32-PGP, BioLegend #101320). The lineage (Lin) antibody cocktail included biotinylated antibodies against FcεR, B220, CD19, Mac-1, Gr-1, CD11c, Ter-119, CD5, CD49b, CD8α, and CD4, followed by staining with fluorophore-conjugated streptavidin. To ensure complete exclusion of NK and T cells from the Lin⁻ gate, cells were routinely stained with antibodies against NK1.1, TCRβ, and TCRδ, each conjugated to distinct fluorophores, and these populations were gated out prior to further analysis. When NK and/or T cells were not stained separately, the corresponding biotinylated antibodies were included in the lineage cocktail. Therefore, Lin^−^ means cells negative all of the markers described above. Flow cytometric analyses and sorting with antibodies against lineage and other markers conjugated with specific fluorophores (Table S2). For intracellular cytokine staining, cells were fixed and stained with antibodies using Cytofix/Cytoperm (BD Bioscience #554714) for 30 min on ice. For intranuclear transcription factor staining, cells were fixed and stained with antibodies using True-Nuclear Transcription Factor Buffer Set (Biolegend #424401), for 45-50 min in room temperature (RT). BrdU incorporation and detection was performed by using BrdU Flow Kits (BD Biosciences #552598). Flow cytometry analyses were performed using an LSR II (BD Biosciences) or an Aurora (Cytek Biosciences) flow cytometer. Data were analyzed by using FlowJo (Version 10.10.0).

Single cell suspensions of human labial salivary gland biopsy tissue were prepared on the same day of biopsy as previously described^59^ and stained with antibodies to CD45, CD3, CD4, and CD8 after blocking with 10% human AB serum and 2% fetal bovine serum in PBS. Non-viable cells were excluded using propidium iodide. Analytical data were collected using a FACSAria (BD Biosciences) sorter.

#### Single-cell RNA sequencing and data analysis

Lin⁻CD127⁺CD45^hi^KLRG1⁻ cells (ILC1s) and Lin⁻TCRδ⁺ (γδ T cells) were sorted from the small-intestinal lamina propria and used for cDNA library preparation according to the manufacturer’s protocol (10x Genomics). Libraries were sequenced on an Illumina NovaSeq 6000 platform.

Cell Ranger pipeline version 3.1.0 (10x Genomics) was used for sample demultiplexing, barcode processing, and single-cell 3′ gene counting. Raw base call (BCL) files generated by the Illumina NovaSeq 6000 were demultiplexed into sample-specific FASTQ files using the cellranger mkfastq command. FASTQ files were then processed using cellranger count for alignment to the mouse reference genome (mm10), filtering, and gene quantification.

The *Seurat* R package (version 4.3) was used for downstream analysis. Canonical correlation analysis was applied for data integration and batch effect correction (Butler et al., 2018; Stuart et al., 2019). Principal component analysis (PCA) was performed on the scaled expression matrix using the 2,000 most variable genes, and the first 20 principal components were used for subsequent analyses. Uniform manifold approximation and projection (UMAP) was performed using these 20 PCs. Cell clustering and re-clustering were conducted using the *FindClusters* function, which implements a shared nearest-neighbor modularity optimization-based algorithm, with resolutions ranging from 0.5 to 1.0 explored.

#### Analysis of TCR gene rearrangements

Desired cell populations were collected using a FACSAria cell sorter and resuspended in proteinase K digestion buffer at a density of 500 cells/µl. Samples were incubated at 55°C for 1 hour, followed by heating at 70°C for 10 minutes. Digested samples were diluted three-fold, and 2 µl of the diluted lysate was used per PCR reaction with primer pairs specific for individual rearrangement events (Table S3). PCR amplification with high-fidelity Taq polymerase (Genesee Scientific, Cat. # 17-108) was performed for 40 cycles, and products were analyzed by agarose gel electrophoresis.

For next-generation sequencing, adaptor sequence was added by using primers listed in Table S4 and initial PCR products as templates. These primers generated shorter PCR products suitable for sequencing. PCR products were purified using KAPA PURE magnetic beads (Fisher Scientific, Cat. # 501965300) to remove free primers. Purified DNA was then used as a template for index (Table S5) primer addition by PCR (8 cycles). Final libraries were sequenced on an Illumina NovaSeq 6000 platform.

Sequencing outputs were processed by using MiXCR software (version 4.4.1) ^60, 61^. Reads from raw data were aligned to the mouse genome in regions between FR3 and FR4 of TCR genes, which include the V, D, and J regions. The alignment output was used to assemble an analysis to identify reads with CDR3 clonotypes based on nucleotide sequences. The results were exported as excel files. The cut-off threshold for clonotype analysis was one clonotype per 10,000 reads. Consensus CDR3 amino acid sequences were generated by using DNASTAR software (version 17.6) with MAFFT algorithm for multiple sequence alignment.

#### Measurement of autoantibodies

Blood was collected from 7–8-month-old Id3-deficient and control mice, and serum samples were frozen in aliquots at −20°C. Autoantibodies were analyzed by indirect immunofluorescence assay using HeLa cells as described previously ^35^. All serum samples were tested at a 1:200 dilution. For ELISA assays, sera were diluted 1:100 and analyzed according to the manufacturers’ instructions: Alpha Diagnostic International Inc. (anti-dsDNA) #5120 and MyBioSource (anti-ANA) #MBS261480.

#### Determination of cytokine production

To measure cytokine production following anti-TCRδ stimulation, Vγ1.1⁺Vδ6.3⁺ T cells were sorted from C57BL/6 and *Id3^f/f^*;*Cd3g*^iCre^ mice and seeded at 20,000 cells per well in antibody-coated 96-well plates. Cells were cultured in complete RPMI-1640 medium (cytiva #SH30011.03) supplemented with 10 ng/ml IL-2 and IL-7 (R&D Systems #402-ML #407-ML). After 3 days of incubation, supernatants were collected and analyzed using Legendplex multiplex assays according to the manufacturer’s instructions (BioLegend #740446 #740819). Data was analyzed using the software provided by the vendor. Serum IL-17 levels were also measured using Legendplex reagents. Initially, both the B-cell and inflammation panels were screened, and no additional cytokines were found to be significantly different between the experimental cohorts.

### QUANTIFICATION AND STATISTICAL ANALYSIS

Data analysis was performed with GraphPad Prism (Version 10.6.892). Data are represented as the mean ± SEM. Normal distribution was determined by using D’Agostino-Pearson omnibus normality test. Analysis of two groups were performed with two-tailed unpaired Mann-Whitney or Welch’s test depending on their distribution. Analysis of three groups or more were performed with Kruskal-Wallis or Ordinary One-Way ANOVA test depending on their distribution. Results were considered statistically significant with *p* < 0.05. Not statistically significant results (*p* ≥ 0.05) were not reported unless indicated as “ns”.

