## supplemental materials; figures and tables for "A feedback control for restraining autoimmune γδ T cells: reprogramming into ILC1s"

**Supplemental figure legends**

**Figure S1. icCD3$^+$ ILC1-like cells in the small intestine do not express TCRδ. (related to Figure 1).**

**a,** Intracellular staining for TCRδ. **b,** icCD3$^+$ ILC1-like cells do not express TCRδ even after 3 days of culture.

**Figure S2. Differential expression of CD8 on γδ T cells (related to Figure 2).**

TCRδ$^+$ cells in the intraepithelial lymphocyte (IEL) and lamina propria (LPL) fractions of small intestine were examined for CD8 expression.

**Figure S3. Analyses of Vγ1.1$^+$Vδ6.3$^+$ cells in the thymus (related to Figure 3)**

**a,** Sorting strategy of cells used in analyses of TCR gene rearrangement shown in Figure 3D. **b,** Expression of PLZF and NK1.1 in Vγ1.1$^+$Vδ6.3$^+$ cells of C57BL/6 mice.

**Figure S4. Analyses of ILC1-like and Vγ1.1$^+$Vδ6.3$^+$ cells in wild type and Id3-deficient mice (related to Figure 4)**

**a,** Analyses of icCD3$^+$ ILC1-like cells in the lamina propria of the small intestine of Id3-deficient and control mice. **b,** The percentages of Vγ1.1$^+$Vδ6.3$^+$ cells in total CD45$^+$ cells in the small intestine of Id3-deficient and control mice are used to assess the levels of these cells.

**Figure S5. Cytokine production by wild type and Id3-deficient Vγ1.1$^+$Vδ6.3$^+$ cells (related to Figure 4).**

**a,** Cytokine levels in the supernatants of $V\gamma1.1^{+}V\delta6.3^{+}$ cells from the spleen of Id3-deficient and control mice cultured in 10 ng/ml IL-2 and IL-7 with or without plate-bound anti-TCRδ for 3 days. **b,** Serum levels of IL-17 in 7-8 months old mice in Id3-deficient and control mice were determined using ELISA. **c,** IL-17 or IFNγ intracellular staining of $V\gamma1.1^{+}V\delta6.3^{+}$ cells that express either RORγT or T-bet after stimulation with PMA and ionomycin for 3 hours *in vitro*.

**Figure S6. Additional analyses of 7-8 months old Id3-deficient and control mice (related to Figure 6).**

**a,** H&E staining of liver sections. **b,** Occasional splenomegaly (~20%) seen in Id3-deficient mice and histology of the spleen sections. **c,** Analyses of T cell compartments in the salivary glands and lung, thymus and spleen. **d,** BrdU incorporation measured for indicated subsets of thymocytes 16 hours after injection of BrdU in 2-month old mice.

Supplemental Figure 1

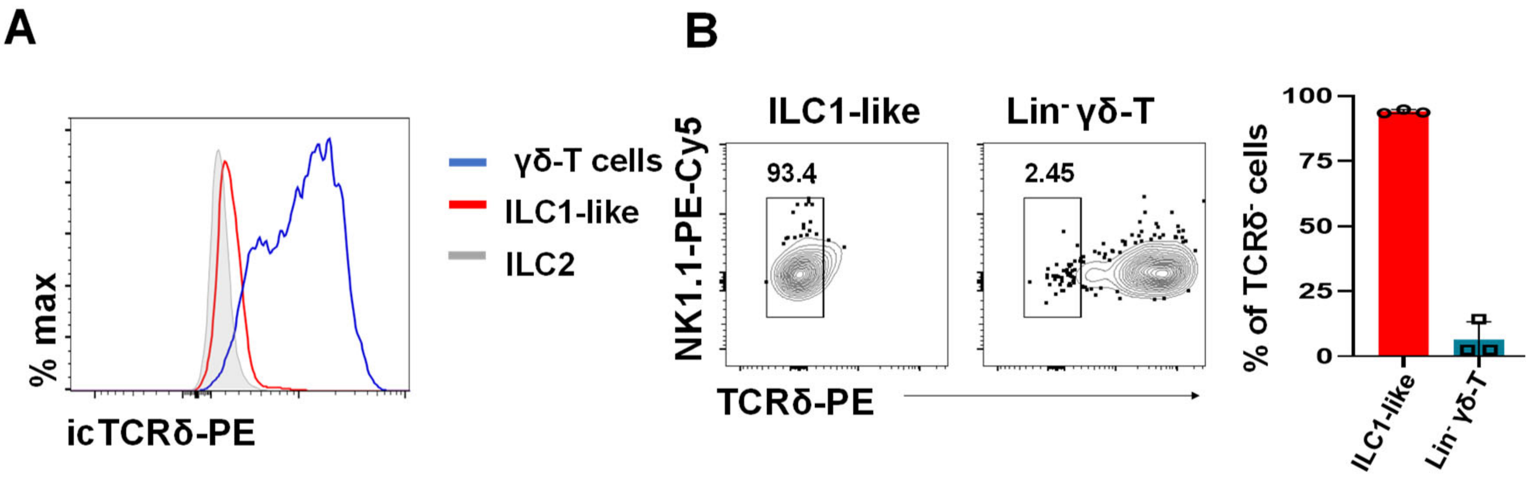

#### Supplemental Figure 2

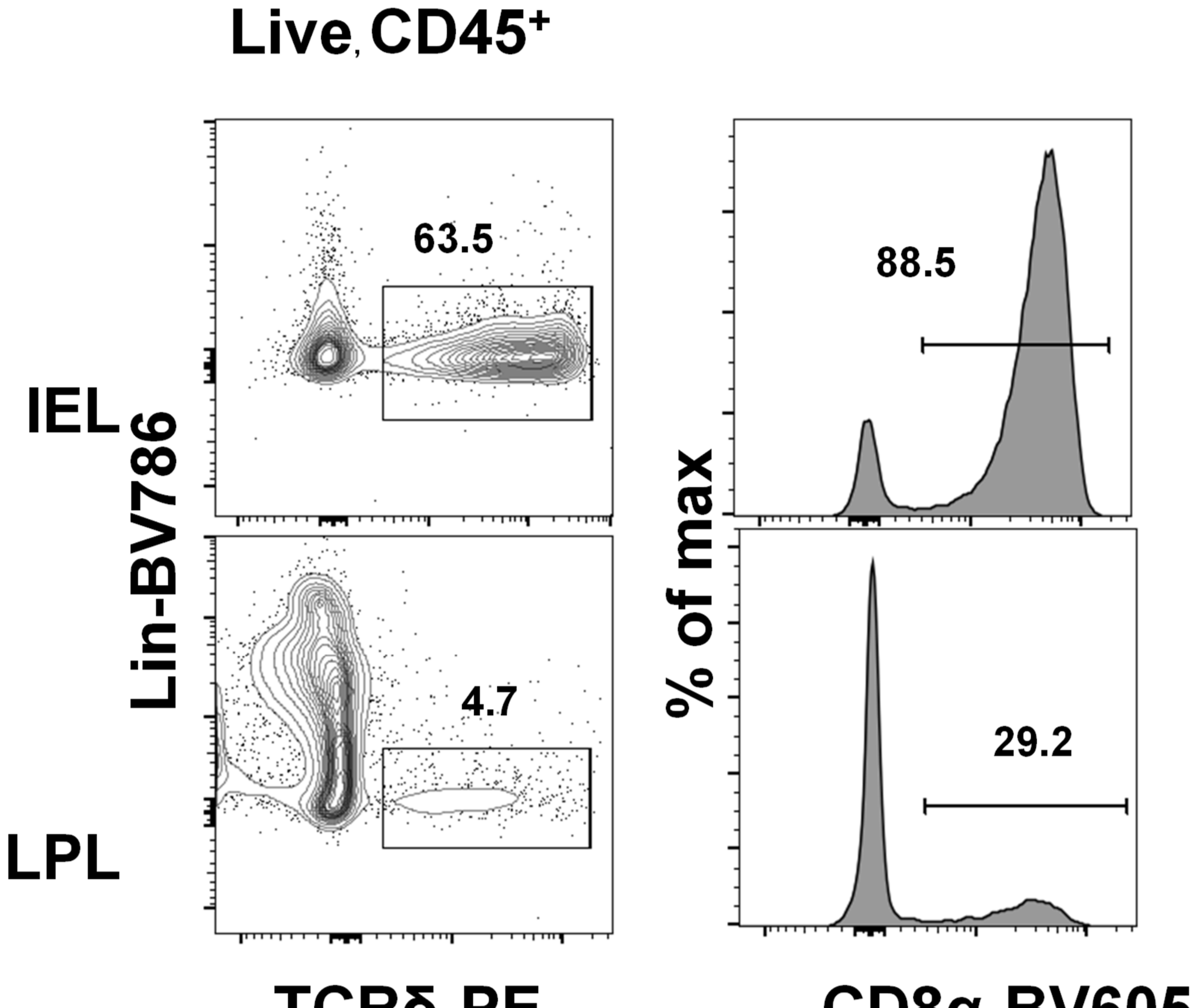

#### Supplemental Figure 3

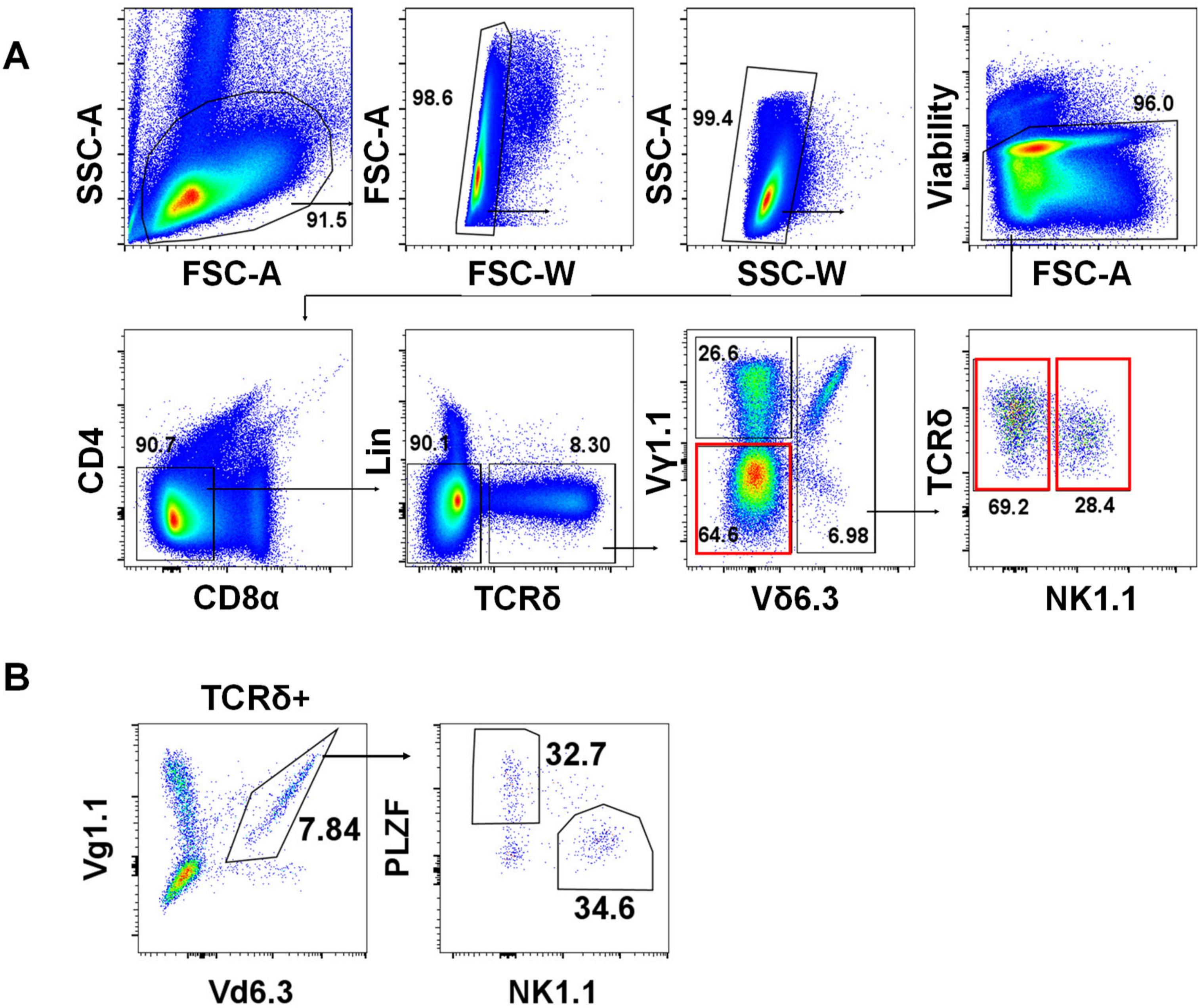

### Supplemental Figure 4

A

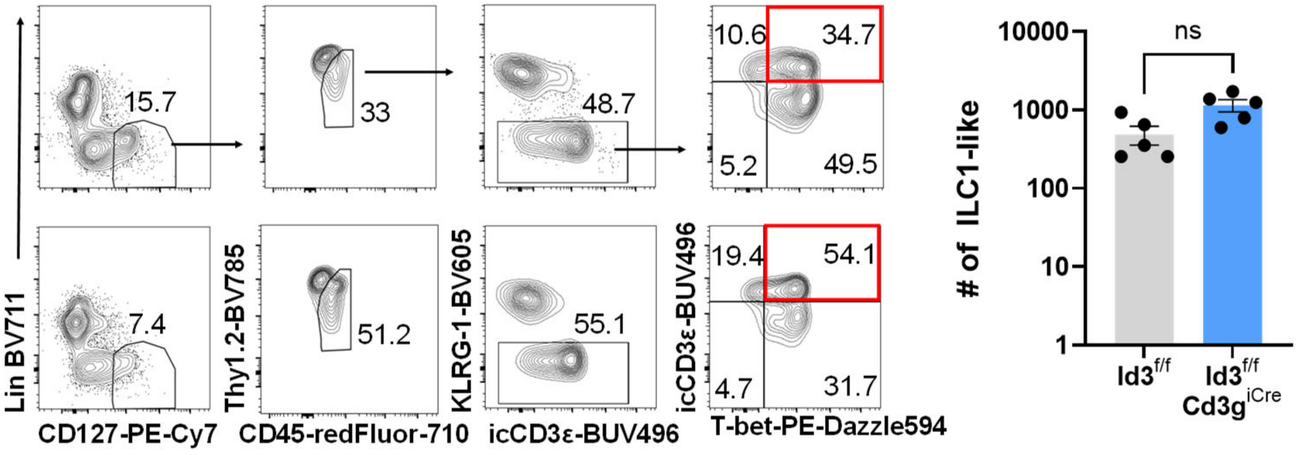


B

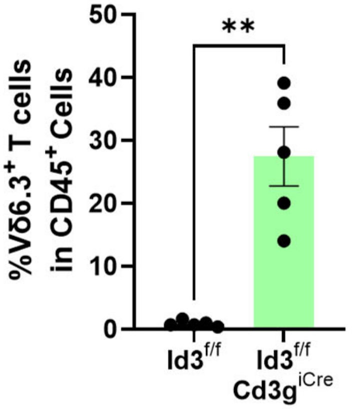

### Supplemental Figure 5

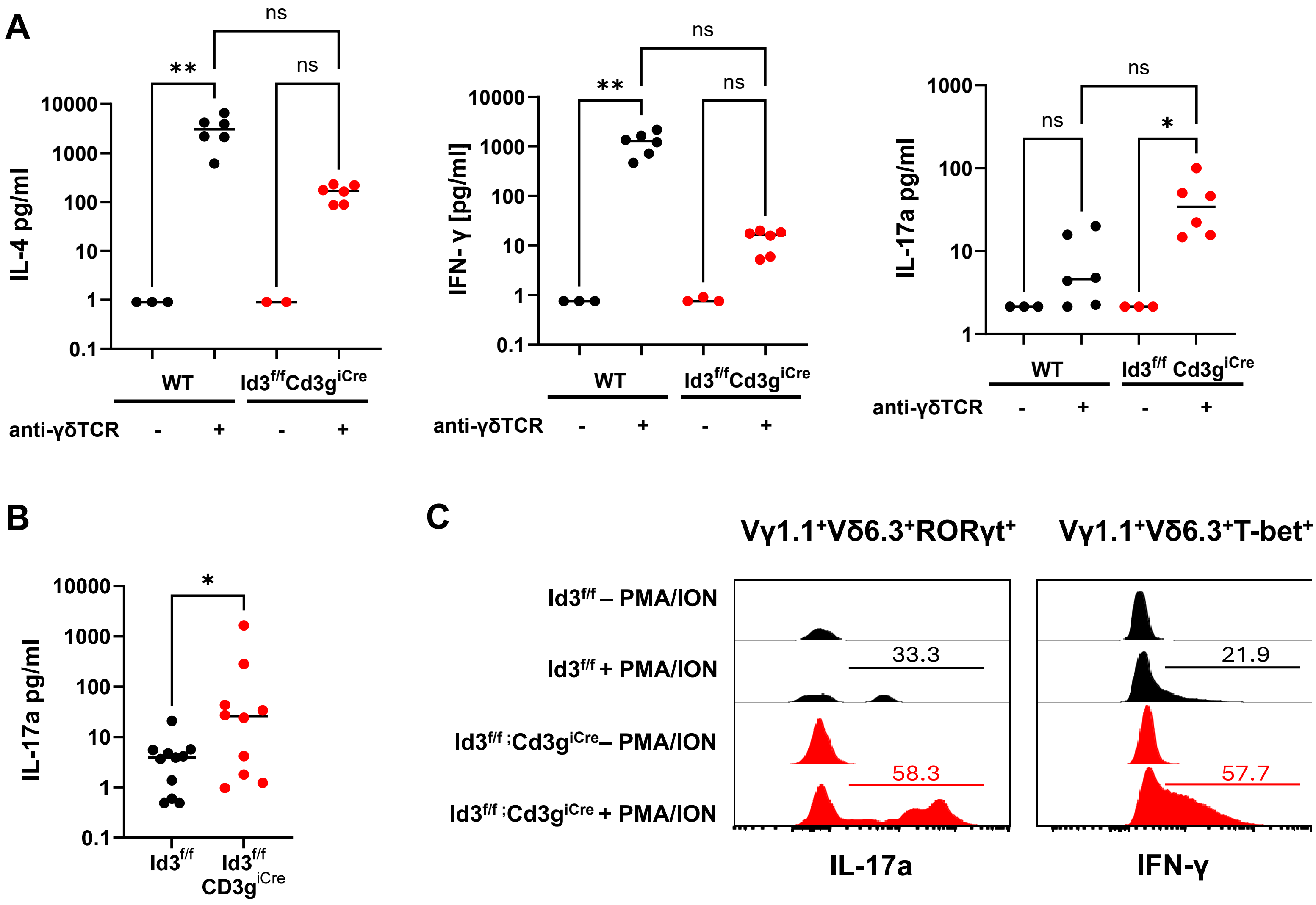

### Supplemental Figure 6

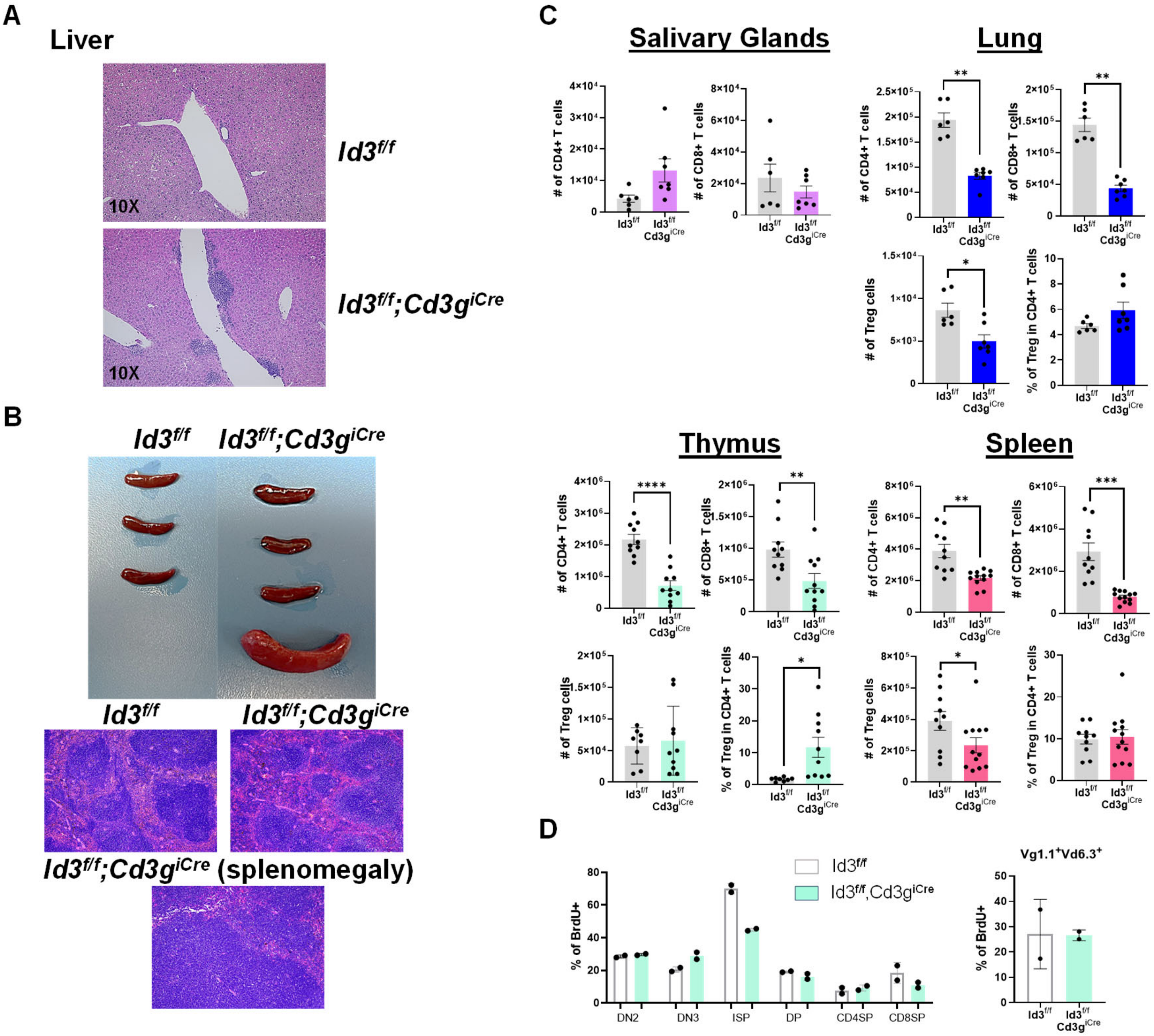

| **Table S1. Demographic and Clinical Features of Human Cohort** | | | | |
|---|---|---|---|---|
| **Feature** | **Ro (+) SjD (n=8)** | **Ro (-) SjD (n=15)** | **Non-SjD (n=38)** | **p-Value** |
| Age (mean (SD)) | 56 (12) | 53 (20) | 49 (14) | 0.438 **a** |
| Race (%) | | | | 0.095 **b** |
| White | 9/15 (60.0) | 8/8 (100) | 16/38 (42.1) | |
| More than one | 6/15 (40) | 0/8 (0) | 17/38 (44.7) | |
| AI/AK Native | 0/15(0) | 0/8(0) | 3/38(7.9) | |
| Black | 0/15 (0) | 0/8 (0) | 2/38 (5.3) | |
| Sex (%) | | | | 0.596 **c** |
| Female | 13/15 (86.7) | 8/8 (100) | 36/38 (94.7) | |
| Clinical Features (%) **d** | | | | |
| FS + | 11/15 (73.3) | 8/8 (100) | 2/15 (13.3) | |
| Ro + | 15/15 (100) | 0/8(0) | 4/38 (10.5) | |
| WUSF + | 10/15 (66.7) | 8/8 (100) | 15/38 (39.5) | |
| Schirmer + | 8/15 (53.3) | 1/8(12.5) | 11/38 (28.9) | |
| OSS + | 8/15 (53.3) | 4/8 (50.0) | 12/38 (31.6) | |
| ESSDAI (%) **e** | | | | |
| Low (0-4) | 4/15 (26.7) | 4/8 (50.0) | N/A | |
| High (≥5) | 11/15 (73.3) | 4/8 (50.0) | N/A | |

**a** One-way ANOVA; **b** Chi-square test; **c** Fisher's exact test; **d** % with positive tests, FS= salivary gland biopsy focus score, Ro=serum anti-Ro/SS-A IgG, WUSF=whole unstimulated salivary flow rate; Schirmer= tear flow rate; OSS=ocular staining score; **e** ESSDAI=European Alliance of Associations for Rheumatology Sjögren's syndrome disease activity index; N/A=not applicable

#### Table S2. List of antibodies used in experiments for flow cytometry and sorting staining

| Specificity | Company/Source | Clone/Identifier |
|---|---|---|
| Syrian hamster anti-Mo KLRG1 PerCP-eFluor710 | Thermo Fisher Scientific | Cat# 46-5893-82, clone 2F1, RRID:AB_10670282 |
| Syrian hamster anti-mouse/human KLRG1 (MAFA) Brilliant Violet 605 | BioLegend | Cat# 138419, clone 2F1/KLRG1 RRID:AB_2563357 |
| Rat anti-mouse CD127 (IL-7Ralpha) PE-Cyanine7 | Thermo Fisher Scientific | Cat# 25-1271-82, clone A7R34, RRID:AB_469649 |
| Rat anti-mouse CD127 (IL-7Ralpha) PE/Dazzle 594 | BioLegend | Cat# 135032, clone A7R34, RRID:AB_2564217 |
| Armenian hamster anti-mouse CD3ε BUV496 | BD Biosciences | Cat# 612955, clone 145-2C11, RRID:AB_2870231 |
| Armenian hamster anti-mouse CD3ε APC | BioLegend | Cat# 100312, clone 145-2C11, RRID:AB_312677 |
| Rat anti-mouse CD90.2 Brilliant Violet 785 | BioLegend | Cat# 105331, clone 30-H12, RRID:AB_2562900 |
| Rat anti-mouse CD90.2 (Thy1.2) APC/Fire 750 | BioLegend | Cat# 140326, clone 53-2.1, RRID:AB_2650963 |
| Rat anti-mouse CD90.2 antibody BUV395 | BD Biosciences | Cat# 565257 , clone 53-2.1, RRID:AB_2739136 |
| Mouse anti-mouse NK-1.1 PE-Cyanine7 | BioLegend | Cat# 108716, clone PK136, RRID:AB_493590 |
| Mouse anti-mouse NK-1.1 Alexa Fluor 647 | BioLegend | Cat# 108720, clone PK136, RRID:AB_2132713 |
| Armenian hamster anti-mouse TCR gamma/delta APC | BioLegend | Cat# 118116, clone GL3, RRID:AB_1731813 |
| Armenian hamster anti-mouse TCR gamma/delta PE | BioLegend | Cat# 118108, clone GL3, RRID:AB_313832 |
| Armenian hamster anti-mouse TCR gamma/delta APC/Fire 750 | BioLegend | Cat# 118136, clone GL3, RRID:AB_2650828 |
| Armenian hamster anti-mouse TCR beta chain Brilliant Violet BV421 | BioLegend | Cat# 109230, clone H57-597, RRID:AB_2562562 |
| Armenian hamster anti-mouse TCR beta chain BUV737 | BD Biosciences | Cat# 612821, clone H57-597, RRID:AB_2870145 |
| Streptavidin Brilliant Violet 711 | BD Biosciences | Cat# 563262, RRID:AB_2869478 |
| Rat anti-mouse CD45 Brilliant Violet BV786 | BD Biosciences | Cat# 564225, clone 30-F11, RRID: AB_2716861 |
| Mouse anti-mouse CD45.2 RedFluor 710 | Tonbo Biosciences | Cat# 80-0454, clone 104, RRID:AB_2621988 |
| Armenian hamster anti-human/mouse/rat CD278 (ICOS) Brilliant Violet BV785 | BioLegend | Cat# 313533, clone C398.4A, RRID:AB_2629728 |
| Armenian hamster anti-human/mouse/rat CD278 (ICOS) PE-Cyanine7 | BioLegend | Cat# 313520, clone C398.4A, RRID:AB_10643411 |
| Rat anti-mouse CD25 Brilliant Violet 785 | BioLegend | Cat# 102051, clone PC61, RRID:AB_2564131 |
| Rat anti-mouse CD25 PE-Cyanine7 | BioLegend | Cat# 102016, clone PC61, RRID:AB_312865 |
| Rat anti-mouse CD8a Brilliant Violet B605 | BioLegend | Cat# 100744, clone 53-6.7, RRID:AB_2562609 |
| Rat anti-mouse CD8a PE-Cyanine7 | BioLegend | Cat# 100722, clone 53-6.7, RRID:AB_312761 |
| Rat anti-mouse CD4 PE-Dazzle | BioLegend | Cat# 100456, clone GK1.5, RRID:AB_2565845 |

| | | |
|---|---|---|
| Rat anti-mouse CD4 APC-Cyanine7 | BioLegend | Cat# 100414, clone GK1.5, RRID:AB_312699 |
| Rat anti-mouse/human CD11b Brilliant Violet 785 | BioLegend | Cat# 101243, clone M1/70, RRID:AB_2561373 |
| Armenian hamster anti-mouse CD11c PerCP | BioLegend | Cat# 117326, clone N418, RRID:AB_2129643 |
| Rat biotin anti-mouse/human CD45R/B220 | BioLegend | Cat# 103204, clone RA3-6B2, RRID:AB_312989 |
| Rat biotin anti-mouse/human CD11b antibody | BioLegend | Cat# 101204, clone M1/70, RRID:AB_312787 |
| Armenian hamster biotin anti-mouse CD11c | BioLegend | Cat# 117304, clone N418, RRID:AB_313773 |
| Armenian hamster biotin FcεRIα | BioLegend | Cat# 134304, clone MAR-1, RRID:AB_1626106 |
| Rat biotin anti-mouse TER-119/Erythroid Cells | BioLegend | Cat# 116204, clone Ter-119, RRID:AB_313705 |
| Mouse biotin anti-mouse CD19 | BioLegend | Cat# 101504, clone MB19-1, RRID:AB_312823 |
| Rat biotinylated anti-mouse CD49b/Pan- NK cells | BD Biosciences | Cat# 553856,clone DX5, RRID:AB_395092 |
| Armenian hamster biotin anti-mouse CD3ε | BioLegend | Cat# 100304,clone 145-2C11, RRID: AB_312669 |
| Rat TruStain FcX (anti-mouse CD16/32) | BioLegend | Cat# 101320, clone 93, RRID:AB_1574975 |
| Mouse anti human/mouse GATA-3 BUV395 | BD Biosciences | Cat# 565448, clone L50-823, RRID: AB_2739241 |
| Mouse anti-mouse RORγt Brilliant Violet BV421 | BD Biosciences | Cat# 562894, clone Q31-378, RRID:AB_2687545 |
| Mouse anti-mouse T-bet PE/Dazzle 594 | BioLegend | Cat# 644828, clone 4B10, RRID:AB_2565677 |
| Rat anti-mouse EOMES PerCP-eFluor 710 | Thermo Fisher Scientific | Cat# 46-4875-82, clone Dan11mag, RRID:AB_10597455 |
| Rat anti-mouse/human CD45R/B220 APC-Fire810 | BioLegend | Cat# 103278, clone RA3-6B2, RRID:AB_2860603 |
| Rat anti-mouse CD19 PE-Cyanine7 | BioLegend | Cat# 115510, clone 6D5, RRID:AB_313645 |
| Rat anti-mouse CD23 PE | BioLegend | Cat# 101607, clone B3B4, RRID:AB_312832 |
| Rat anti-mouse CD21 Pacific Blue | BioLegend | Cat# 123414, clone 7E9, RRID:AB_2085158 |
| Rat anti-mouse/human GL7 Alexa Fluor F647 | BioLegend | Cat# 144606, clone GL7, RRID:AB_2562185 |
| Rat anti-mouse CD38 PE-Cyanine7 | BioLegend | Cat# 102718, clone 90, RRID:AB_2275531 |
| Armenian hamster CD279 (PD-1) APC-eFluor780 | Thermo Fisher Scientific | Cat# 47-9985-80, clone J43, RRID:AB_2574002 |
| Rat anti-mouse CD279 (PD-1) APC-Fire810 | BioLegend | Cat# 135252, clone 29F.1A12, RRID:AB_2910292 |
| Rat anti-mouse CD185 (CXCR5) Brilliant Violet 650 | BioLegend | Cat# 145517, clone L138D7, RRID:AB_2562453 |
| Rat FOXP3 APC | Thermo Fisher Scientific | Cat# 17-5773-82, clone FJK-16s, RRID:AB_469457 |
| Armenian hamster anti-mouse TCR Vγ1.1 Pacific Blue | BioLegend | Cat# 141110, clone 2.11, RRID:AB_2750499 |
| Rat anti-mouse TCR Vδ6.3 APC | Biolegend | Cat# 154806, clone C504.17C, RRID:AB_2728221 |
| Rat anti-mouse TCR Vδ6.3 PE | BioLegend | Cat# 154803, clone C504.17C, RRID:AB_2728219 |

| | | |
|---|---|---|
| Mouse anti-mouse/human PLZF Alexa Fluor 488 | Thermo Fisher Scientific | Cat# 53-9320-82, clone Mags.21F7, RRID:AB_2574445 |
| Mouse anti-mouse/human PLZF PE | Thermo Fisher Scientific | Cat# 12-9320-82, clone Mags.21F7, RRID:AB_11148934 |
| Mouse anti-mouse/human Bcl-6 PE-CF594 | BD Biosciences | Cat# 562401, clone K112-91, RRID:AB_11152084 |

#### Table S3. Primer sequences for PCR rearrangements.

| Gene | Forward primer | Reverse primer |
|---|---|---|
| *Vg1.1-Jg4* | 5'-GAGAGTGCGCAAATATCCTGTTATA | 5'-TGGGGGAATTACTACGAGCT |
| *Vg1.2-Jg2* | 5'-CTTCCATATTTCTCCAACACAGC | 5'-ACTATGAGCTTTGTTCCTTCTGCAA |
| *Vg2-Jg1* | 5'-TGGACATGGGAAGTTGGAG | 5'-CAGAGGGAATTACTATGAGC |
| *Vg3-Jg1* | 5'-GATCAGCTCTCCTTTACCC | 5'- CAGAGGGAATTACTATGAGC |
| *Vg4-Jg1* | 5'-CTGGGGTCATATGTCATCAA | 5'- CAGAGGGAATTACTATGAGC |
| *Vd4-Jd1* | 5'-CCGCTTCTCTGTGAACTTCC | 5'- CAGTCACTTGGGTTCCTTGTCC |
| *Vd5-Jd1* | 5'-CAGATCCTTCCAGTTCATCC | 5'- CAGTCACTTGGGTTCCTTGTCC |
| *Vd1-Jd1* | 5'-GGTGGAAAGAGCAACCTCAAAG | 5'- CAGTCACTTGGGTTCCTTGTCC |
| *Vd6.3-Jd1* | 5'-CTGGTACAAGCACCTTCTTAGTGGAG | 5'- CAGTCACTTGGGTTCCTTGTCC |
| *Id2* | 5'- | 5'- |

#### Table S4. Adaptor primer sequences.

| Gene | Primer sequence 5' -> 3' |
|---|---|
| *Vg1.1-Jg4* | **F** TCGTCGGCAGCGTCAGATGTGTATAAGAGACAGGATTTTCAAACTTCTACCTCAACC<br>**R** GTCTCGTGGGCTCGGAGATGTGTATAAGAGACAGTGGGGGAATTACTACGAGCT |
| *Vg2-Jg1* | **F** TCGTCGGCAGCGTCAGATGTGTATAAGAGACAGTATATTCCTTGGAGGAAGAAGAC<br>**R** GTCTCGTGGGCTCGGAGATGTGTATAAGAGACAGACCAGAGGGAATTACTATGAGC |
| *Vd4-Jd1* | **F** TCGTCGGCAGCGTCAGATGTGTATAAGAGACAGCCGCTTCTCTGTGAACTTCC<br>**R** GTCTCGTGGGCTCGGAGATGTGTATAAGAGACAGCAGTCACTTGGGTTCCTTGTCC |
| *Vd6.3-Jd1* | **F** TCGTCGGCAGCGTCAGATGTGTATAAGAGACAGCCTTGTCATTTCAACCTTACAACC<br>**R** GTCTCGTGGGCTCGGAGATGTGTATAAGAGACAGCAGTCACTTGGGTTCCTTGTCC |

#### Table S5. Index primer sequences.

| Index 1 | Sequence |
|---|---|
| N701 | CAAGCAGAAGACGGCATACGAGATTAAGGCGAGTCTCGTGGGCTCGG |
| N702 | CAAGCAGAAGACGGCATACGAGATCGTACTAGGTCTCGTGGGCTCGG |
| N703 | CAAGCAGAAGACGGCATACGAGATAGGCAGAAGTCTCGTGGGCTCGG |
| N704 | CAAGCAGAAGACGGCATACGAGATTCCTGAGCGTCTCGTGGGCTCGG |
| N705 | CAAGCAGAAGACGGCATACGAGATGGACTCCTGTCTCGTGGGCTCGG |
| N706 | CAAGCAGAAGACGGCATACGAGATTAGGCATGGTCTCGTGGGCTCGG |
| N707 | CAAGCAGAAGACGGCATACGAGATCTCTCTACGTCTCGTGGGCTCGG |
| N708 | CAAGCAGAAGACGGCATACGAGATCAGAGAGGGTCTCGTGGGCTCGG |
| **Index 2** | **Sequence** |

| | |
|---|---|
| N501 | AATGATACGGCGACCACCGAGATCTACACTAGATCGCTCGTCGGCAGCGTC |
| N502 | AATGATACGGCGACCACCGAGATCTACACCTCTCTATTCGTCGGCAGCGTC |
| S503 | AATGATACGGCGACCACCGAGATCTACACTATCCTCTTCGTCGGCAGCGTC |
| S504 | AATGATACGGCGACCACCGAGATCTACACGTAAGGAGTCGTCGGCAGCGTC |
| S505 | AATGATACGGCGACCACCGAGATCTACACGTAAGGAGTCGTCGGCAGCGTC |
